# USP24 is an ISG15 cross-reactive deubiquitinase that mediates IFN-I production by de-ISGylating the RNA helicase MOV10

**DOI:** 10.1101/2024.09.06.611391

**Authors:** Rishov Mukhopadhyay, Simeon D. Draganov, Jimmy J. L. L. Akkermans, Marjolein Kikkert, Klaus-Peter Knobeloch, Günter Fritz, María Guzmán, Sonia Zuñiga, Robbert. Q. Kim, Benedikt M. Kessler, Adán Pinto-Fernández, Paul P. Geurink, Aysegul Sapmaz

**Affiliations:** Department of Cell and Chemical Biology, Division of Chemical Biology and Drug Discovery, Leiden University Medical Center, Einthovenweg 20, 2333 ZC, Leiden, The Netherlands; Chinese Academy for Medical Sciences Oxford Institute, Nuffield Department of Medicine, University of Oxford, Roosevelt Drive, Oxford OX3 7BN, United Kingdom; Target Discovery Institute, Nuffield Department of Medicine, Centre for Medicines Discovery, University of Oxford, Oxford, Roosevelt Drive, Oxford, OX3 7FZ, United Kingdom; Department of Cell and Chemical Biology and Oncode Institute, Leiden University Medical Center LUMC, Einthovenweg 20, 2333 ZC, Leiden, The Netherlands; Department of Medical Microbiology, Leiden University Medical Center, Molecular Virology Laboratory, Leiden, The Netherlands; Institute of Neuropathology, Faculty of Medicine, Department of Molecular Genetics, University of Freiburg, Freiburg 79106, Germany; and Centre for Integrative Biological Signalling Studies, Department of Molecular Genetics, University of Freiburg, Freiburg 79104, Germany; Department of Cellular Microbiology, University of Hohenheim, Stuttgart 70599, Germany; Department of Molecular and Cell Biology, National Center of Biotechnology (CNB-CSIC), Madrid, Spain

**Keywords:** ISG15, USP24, DUBs, MOV10, cross-reactivity, and immune response

## Abstract

The interferon-stimulated gene 15 (ISG15) is a ubiquitin-like modifier induced by type I Interferon (IFN-I) and plays a crucial role in the innate immune response against viral infections. ISG15 is conjugated to target proteins by an enzymatic cascade through a process called ISGylation. While ubiquitin-specific protease 18 (USP18) is a well-defined deISGylase counteracting ISG15 conjugation, ISG15 cross-reactive deubiquitylating enzymes (DUBs) have also been reported. Our study reports USP24 as a novel ISG15 cross-reactive DUB identified through activity-based protein profiling (ABPP). We demonstrate that recombinant USP24 processed pro-ISG15 and ISG15-linked synthetic substrates *in vitro*. Moreover, the depletion of USP24 significantly increased the accumulation of ISG15 conjugates upon IFN-β stimulation. An extensive proteomic analysis of the USP24-dependent ISGylome, integrating total proteome, GG-peptidome, and ISG15 interactome data, identified the helicase Moloney leukemia virus 10 (MOV10) as a specific target of USP24 for deISGylation. Further validation in cells revealed that ISGylated MOV10 enhances IFN-β production/secretion, whereas USP24 deISGylates MOV10 to negatively regulate the innate immune response. This study showcases USP24’s novel roles in modulating ISGylation and modulation of the IFN-I-dependent immune responses, with potential therapeutic implications in infectious diseases, cancer, autoimmunity, and neuroinflammation.

## INTRODUCTION

Ubiquitin (Ub) and ubiquitin-like proteins (Ubls) constitute a highly important, yet complex class of protein post-translational modifications (PTMs) that regulate a wide variety of cellular mechanisms by maintaining protein stability,^1,2^ protein localization,^3^ and protein-protein interactions.^4^ The interferon-stimulated gene 15 (ISG15), a 17 kDa ubiquitin-like protein consisting of two Ubl domains, ^5^ emerges as a key player in innate immunity and is tightly regulated by type I Interferon (IFN-I) signalling. Upon maturation of endogenously expressed pro-ISG15 by liberating the carboxy-terminal LRLRGG motif, ^6^ ISG15, similar to Ub, is conjugated to target proteins via the conjugation machinery that includes the E1 activating enzyme Ube1L,^7^ the E2 conjugating enzyme Ube2L6,^8^ and the E3 ligases HERC5,^9^ TRIM25,^10^ and HHARI/ARIH1,^11^ in humans.^6,12^ The deconjugation of ISG15 from target proteins and the maturation of ISG15 are established by deISGylases.^13^ The Ubiquitin-specific protease 18 (USP18) does not process Ub, but has been characterized as the canonical deISGylase that removes conjugated ISG15 and matures proISG15, and its role in innate immunity is well-documented.^14,15^ Evidence of interplay between the Ub and ISG15 systems has emerged over the last two decades. For instance, Ube2L6,^16^ TRIM25^10^ and HHARI/ARIH1^11^ are involved in the conjugation of both Ub and ISG15. In addition, previous *in vitro* studies using ISG15 activity-based probes identified ISG15 cross-reactivity of a subset of deubiquitinating enzymes (DUBs), including USP2, USP5, USP13, USP14, and USP21.^17,18,19^ We and others have recently unveiled USP16 and USP36 as additional ISG15 cross-reactive DUBs that are able to deconjugate as well as mature ISG15, and we highlighted the potential role of USP16-dependent deISGylation in metabolic processes.^20, 21^ Building upon these discoveries, we aimed to explore additional DUBs with ISG15 cross-reactive properties that may exist, thereby enhancing our understanding of ISG15-mediated cellular processes.

In this study, ISG15 activity-based probes and quantitative activity proteomics were employed to selectively pull-down and identify deISGylating enzymes in murine EL4 cell extracts. Through this approach, USP24 emerged as a novel ISG15 cross-reactive deubiquitinating enzyme (DUB) alongside previously documented cross-reactive DUBs, such as USP5, USP16, and USP14. USP24 is a cysteine protease with a molecular weight of 294 kDa, which was shown to deubiquitinate a variety of substrates, including DDB2 (regulating DNA damage response),^22^ p300 (affecting IL6 expression),^22^ βTrCP (in lung cancer cells),^22^ and p53 (modulating apoptosis).^23^ Our findings revealed that USP24 effectively cleaves pro-ISG15 and ISG15-based substrates *in vitro*, and its depletion results in an increase in cellular ISG15 conjugates. Further, we delineated specific ISGylated targets of USP24 by combining the data sourced from the total proteome, ISGylome, and GG-peptidome analyses. This comprehensive approach was performed in USP24 knockdown (KD) compared to control HeLa (human cervix cancer) cells after treatment with type I Interferon (IFN-I). We discovered that the RNA helicase Moloney leukemia virus 10 protein (MOV10), reported to provide antiviral responses against RNA viruses by enhancing Type I Interferon response, ^24,25^ was explicitly targeted by USP24 for deISGylation, but not the main deISGylase, USP18. Further investigation indicated that depletion of USP24 leads to increased secretion of IFN-β, aligning with previous research findings in viral infection systems,^26^ depending on ISGylation of MOV10 in Huh7 cells. Collectively, our findings propose an ISG15-mediated functional control of MOV10 in the IFN-I mediated immune response controlled by the deISGylase activity of USP24, implicating the potential of USP24 as a valuable therapeutic target with potential applications in infectious diseases, cancer, autoimmunity, and neuroinflammation.

## RESULTS

### Discovery of USP24 as an ISG15-reactive DUB by ABPP

To identify and confirm ISG15 cross-reactive DUBs, we employed activity-based probes equipped with a propargylamide warhead that can form a covalent linkage with the active site cysteine within the catalytic pocket of DUBs. The reactivity of an enzyme with a probe was assessed through either a fluorescent or biotin tag attached to the N-terminus of the probe (Figure 1A).^27^ Initially, we conducted an activity-based pull-down assay using mouse EL4 cell lysates, which were treated with either DMSO or a biotin-tagged full-length mouse ISG15 propargylamide probe (Biotin-mISG15FL-PA).^13^ Probe-bound enzymes were enriched using Neutravidin beads, washed under denaturing conditions, separated via SDS-PAGE, and subsequently subjected to in-gel trypsin digestion and label-free LC-MS/MS analysis. This not only confirmed previously reported ISG15 cross-reactive DUBs such as USP5, USP14, and USP16 but also unveiled USP24 as a so-far unidentified ISG15 cross-reactive DUB (Figure 1B).

**Figure 1:**
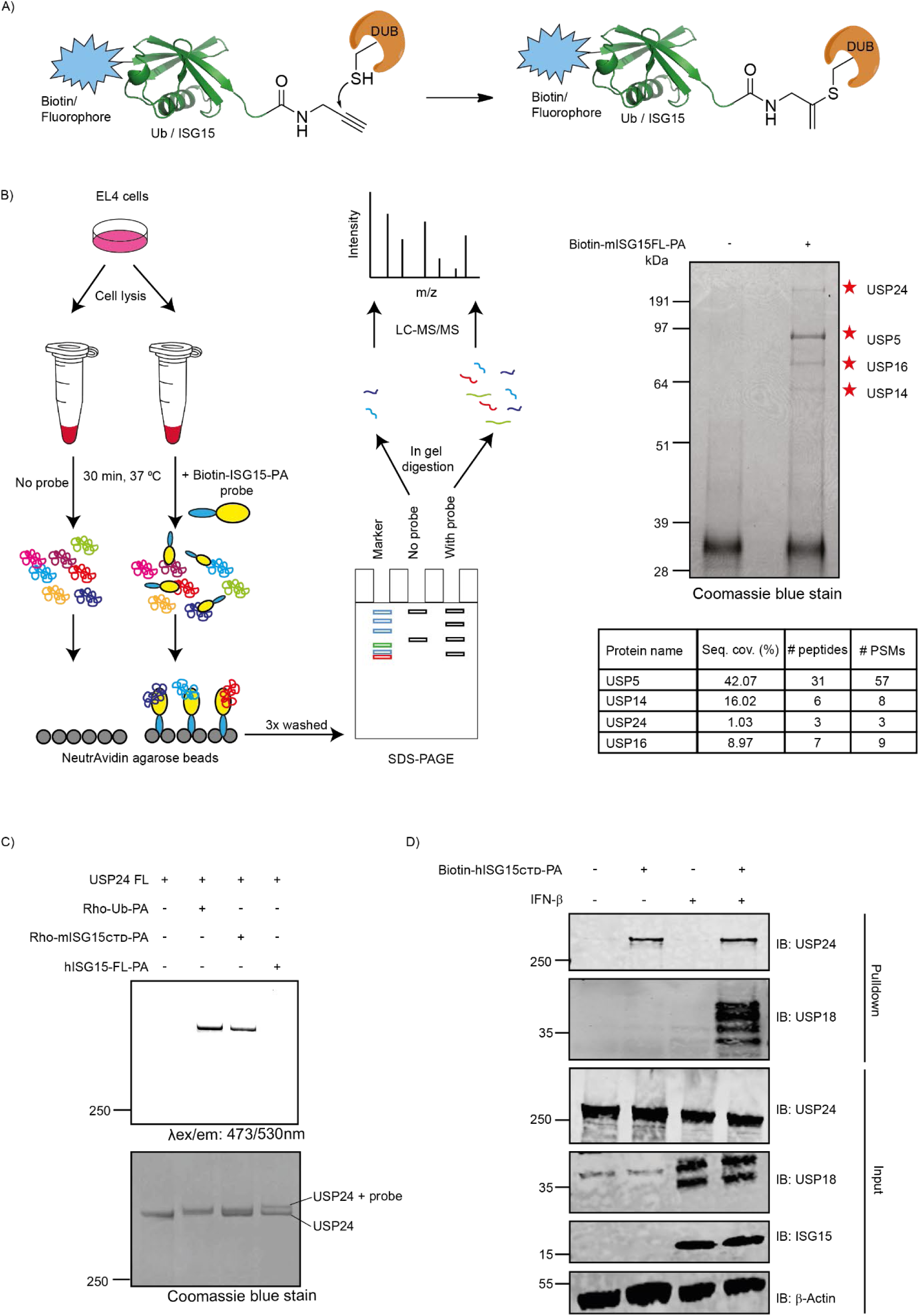
Discovery of USP24 as an ISG15-reactive DUB by ABPP. **(A)** Schematic representation of the covalent modification of a cysteine DUB by Ub/Ubl probes with a propargylamide (PA) warhead. **(B)** Schematic representation of the activity-based pulldown experiment followed by LC-MS/MS analysis. Pulldown and identification of ISG15 cross-reactive DUBs. EL4 cell lysate was incubated with 5 µM of Biotin-mISG15-PA probe or DMSO for 1 h at 37°C, followed by pulldown with Neutravidin beads, separation on SDS-PAGE gel, and coomassie staining. The bands observed in the gel were subjected to LC-MS/MS. ISG15 cross-reactive DUBs identified in mass spectrometry analysis are indicated with red asterisks. Data from mass spectrometry analysis is provided in the table below. **(C)** *In vitro* labeling of recombinant USP24FL protein with fluorescent Rho-Ub-PA and Rho-mISG15_CTD_-PA probes and non-fluorescent hISG15FL-PA probe. USP24FL (0.5 µM) was incubated with 5 µM of indicated probes for 1 h at 37°C, followed by a fluorescence scan (top panel) and Coomassie staining (bottom panel) showing the fraction of bound/unbound USP24. **(D)** Confirmation of endogenous USP24 pull-down with Biotin-hISG15_CTD_-PA probe. HeLa cell lysates from IFN-β stimulated or untreated cells were incubated with 5 µM of Biotin-hISG15_CTD_-PA or DMSO for 1 h at 37°C, followed by pulldown with Neutravidin beads and immunoblot analysis against USP24, along with USP18 and ISG15 as positive controls for IFN-β stimulation and β-actin as loading control. See also Supplementary Fig. 1.

To validate the cross-reactivity of USP24 with ISG15 probe *in vitro*, recombinant full-length human USP24 (USP24FL) was incubated with three fluorescent probes, including Rhodamine-tagged Ub propargylamide (Rho-Ub-PA),^28^ mouse C-terminal domain ISG15 propargylamide (Rho-mISG15_CTD_-PA),^29^ and untagged human full-length-ISG15 propargylamide (hISG15FL-PA) probes.^13^ Labeling of USP24 with Rho-mISG15_CTD_-PA was observed in both fluorescence scanning and coomassie staining, while a clear band shift of recombinant USP24 upon incubation with an untagged hISG15FL-PA probe was observed in coomassie staining, confirming the reactivity of recombinant USP24FL with ISG15 probes (Figure 1C). To further demonstrate the cross-reactivity of cellular USP24 with ISG15, we conducted a pull-down experiment where we incubated lysates from both Interferon-β (IFN-β) stimulated and non-stimulated HeLa cell extracts with Biotin-hISG15_CTD_-PA probe, followed by the enrichment of the probe-bound DUBs with Neutravidin beads. Subsequent analysis by western blot showed that endogenous USP24 was remarkably enriched by the Biotin-hISG15ct-PA probe in lysates from both IFN-β stimulated and non-stimulated cell extracts (Figure 1D), consistent with data obtained from LC-MS/MS analysis. Notably, USP18 was solely enriched in IFN-β stimulated cell lysates, aligning with its known status as an interferon-stimulated gene. Similar results were obtained in HAP1 cells (Supplementary Fig. 1). Taken together, these findings strongly support the conclusion that deubiquitinase USP24 exhibits cross-reactivity with ISG15.

### Characterization of USP24 as an ISG15 processing enzyme and deISGylase

To assess the catalytic capability of USP24 as deISGylase, we performed a series of *in vitro* experiments using ISG15-based substrates. ISG15 is synthesized as an immature protein, requiring the cleavage of a small c-terminal peptide by USP18 or other deISGylating enzymes.^30^ Therefore, we first investigated whether USP24 can process the pro-ISG15 precursor (a.a. 1-165) into mature ISG15 (a.a. 1-157) (Figure 2A, B). Both USP24FL and USP24 catalytic domain (USP24CD) were incubated with recombinant pro-ISG15, and samples were analyzed over multiple time points by SDS-PAGE and coomassie staining. Recombinant USP18 was included as a positive control. Results showed that USP24FL cleaved pro-ISG15 *in vitro*, albeit with lower efficiency compared to USP18. Notably, USP24CD showed no activity in processing pro-ISG15, suggesting the involvement of additional domains (Figure 2C). We further investigated the deISGylase activity of USP24 using an ISG15_CTD_-based substrate that mimics the native isopeptide linkage between ISG15 and the lysine ε-amine of a substrate protein. In this substrate, ISG15_CTD_ is linked to the lysine side chain of a fluorescent TAMRA-Lys-Gly peptide (TAMRA-Lys(ISG15_CTD_)-Gly), and its protease-mediated cleavage can be monitored by fluorescence polarization (FP) (Figure 2D). This revealed that USP24FL can also process this isopeptide-linked ISG15 substrate, although at higher enzyme concentrations than required for processing of the isopeptide-linked Ub subsrate (Figure 2E,F). Notably, USP24CD showed a significant overall decrease in activity towards the Ub substrate, as compared to full-length USP24, requiring about 125 times higher enzyme concentration, and it was hardly active on the ISG15 substrate (Figure 2G-H). We also validated the deISGylase activity of USP24 using fluorogenic substrates ubiquitin rhodamine-morpholine (Ub-Rho-MP) and ISG15 C-terminal domain rhodamine-morpholine (ISG15 _CTD_-Rho-MP) (Supplementary Fig. 2A). USP24FL cleaved both ISG15 and ubiquitin substrates though its activity towards ubiquitin was again higher than towards ISG15 (Supplementary Fig. 2B, C). As expected, USP24 also showed lower activity towards ISG15 compared to USP18 in this *in vitro setting* (Supplementary Fig. 2D). On the other hand, USP24CD processed the ubiquitin substrate but not the ISG15 substrate exhibiting consistent results with the FP assay, suggesting the essentiality of the remaining domains of USP24 for its deISGylase activity (Supplementary Fig. 2E, F). In summary, USP24FL functions as both a pro-ISG15 processing enzyme and a deISGylase, but not its isolated catalytic domain.

**Figure 2:**
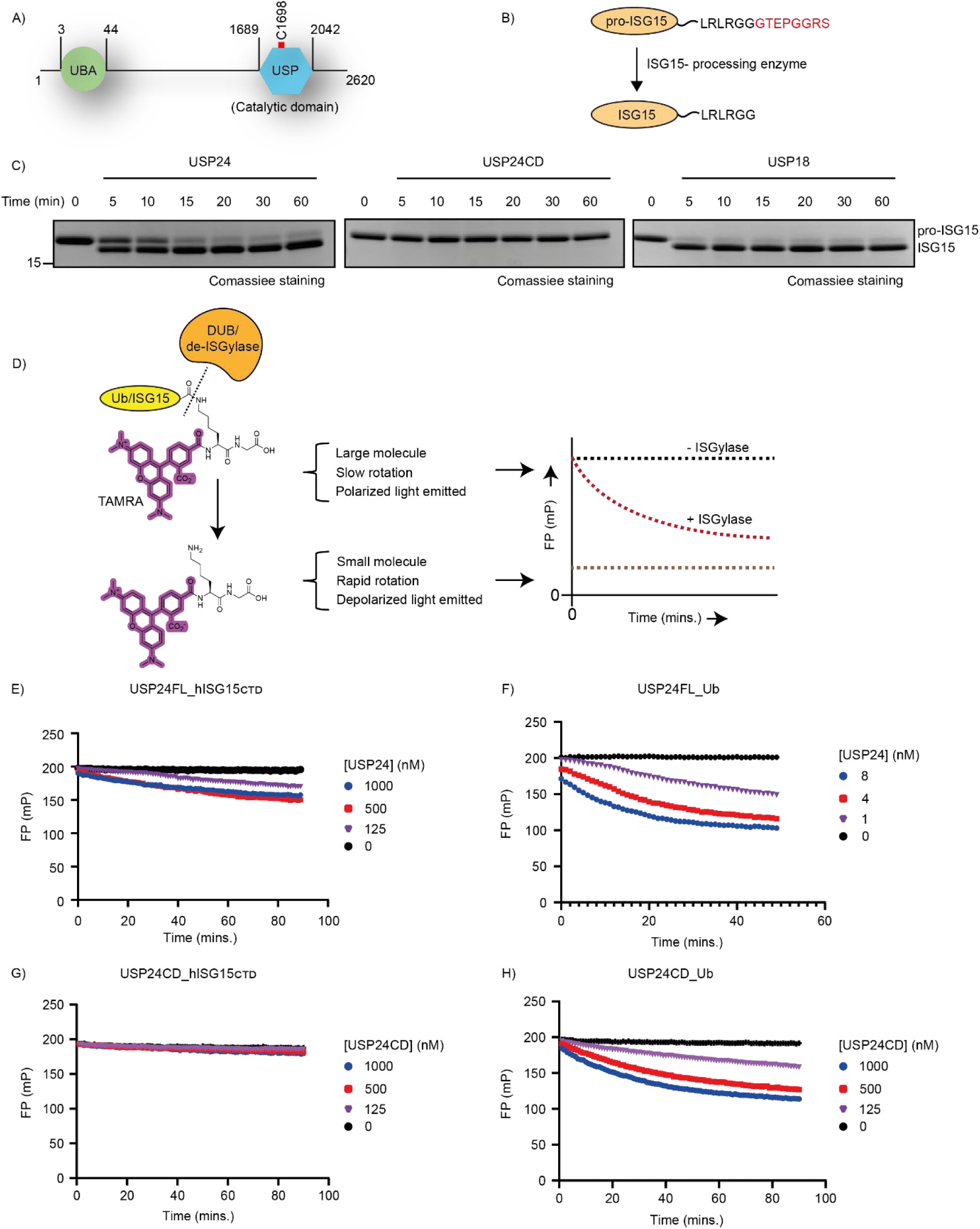
USP24FL but not USP24CD processes ISG15 substrates in vitro. **(A) Schematic representation of USP24FL showing the UBA domain at the N-terminal and catalytic domain (active cysteine 1698 in red) towards the C-terminal end. (B)** Schematic representation of the maturation of precursor ISG15 to mature ISG15 by ISG15 processing enzyme. **(C)** The maturation of recombinant pro-ISG15 by recombinant USP24. 5 µM final concentration of pro-ISG15 was incubated with recombinant USP24FL and USP24CD (0.5 µM) for indicated time points, followed by SDS-PAGE and Coomassie staining. USP18 is used as a positive control. **(D)** Schematic representation of the principle of fluorescence polarization assay to monitor the activity of deconjugating enzymes. The catalytic activity of **(E)** recombinant human USP24FL towards isopeptide-linked human TAMRA-Lys(ISG15_CTD_)-Gly substrate, **(F)** recombinant human USP24FL towards isopeptide-linked human TAMRA-Lys(Ub)-Gly substrate, **(G)** recombinant human USP24CD towards isopeptide-linked human TAMRA-Lys(ISG15_CTD_)-Gly substrate, and **(H)** recombinant human USP24CD towards isopeptide-linked human TAMRA-Lys(Ub)-Gly substrate. Indicated protein concentration was incubated with 200 nM substrate and measured for 90 mins. Substrate cleavage was monitored on the basis of the change in fluorescence polarization (in millipolarization units (mP)). See also Supplementary Fig. 2.

### USP24 depletion increases cellular ISGylation without directly impacting type I IFN signaling

As USP24 appears to cleave pro-ISG15 and ISG15 substrates *in vitro*, we proceeded to investigate its potential deISGylation function in human cells. A time course experiment in HeLa cells was conducted to determine the optimal time for ISG15 conjugation upon IFN-β stimulation, revealing a peak in ISG15 expression and conjugation after 48 hours (Supplementary Fig. 3A,B). Notably, unlike USP18 and ISG15, the levels of USP24 remained unaffected by IFN-β stimulation, suggesting that USP24 is not an interferon-stimulated gene. USP24 siRNA depletion led to a significant accumulation of ISG15 conjugates in HeLa cells upon IFN-β stimulation (Figure 3A), consistent with observations from USP24 knockout (USP24KO) HeLa cells (Supplementary Fig. 3C).

**Figure 3:**
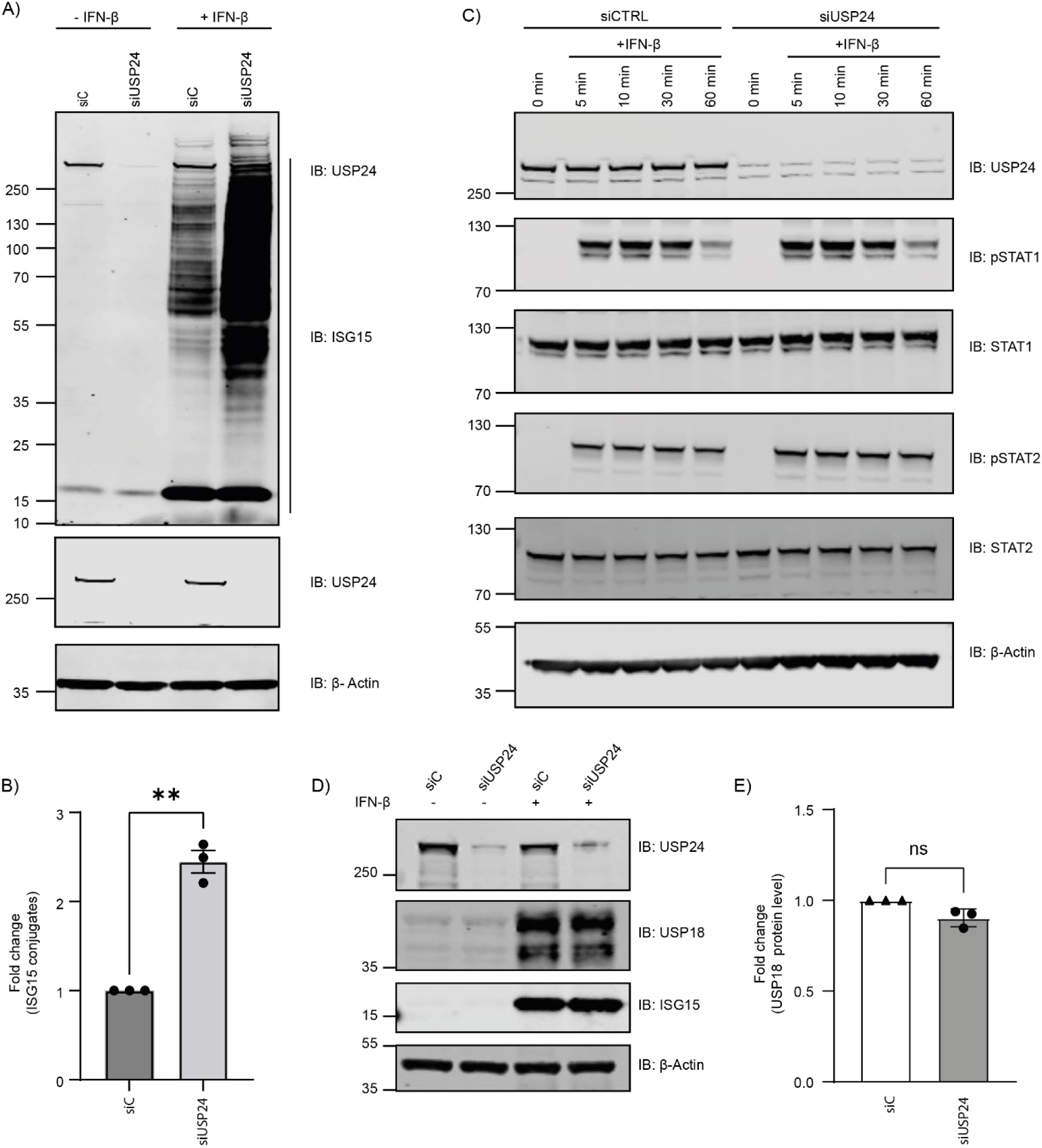
USP24 depletion elevates cellular ISG15 conjugates without impacting the type I IFN signaling cascade. **(A)** Immunoblot of ISG15 conjugation in response to depletion of USP24 in HeLa cells with and without IFN-β stimulation. See also Supplementary Fig. 3C. **(B)** Quantification of the fold change of ISG15 conjugation in IFN-β stimulated HeLa cells as a result of USP24 depletion, n=3 independent experiments. Samples were first normalized as per β-actin followed by normalizing with siC + IFN-β treated sample. Bar graphs report mean, error bars reflect ± s.d. All significant values were calculated using Student’s t-test: **p < 0.05. **(C)** Effect of USP24 depletion on Type I Interferon pathway. HeLa cells transfected with siC or siUSP24 were stimulated with IFN-β for the indicated times. Immunoblot of total STAT1 and STAT2 proteins and phosphorylated STAT1 and STAT2 is provided along with immunoblots of USP24 to prove efficient knockdown and β-actin as loading control. **(D)** Immunoblot to confirm unchanged protein levels of USP18 upon USP24 depletion. HeLa cells were depleted of USP24 using USP24-specific siRNA along with IFN-β treatment for 48 h. Cell lysates from both IFN-β treated and untreated cells were subjected to SDS-PAGE. β-Actin served as loading control. **(E)** Quantification of USP18 protein levels upon USP24 depletion in HeLa cells (n=3) following IFN-β treatment. Data was normalized to siC. Bar graphs report mean, error bars reflect ± s.d. All significant values were calculated using Student’s t-test: ns = not significant.

To exclude any indirect effects of USP24 depletion on Type I IFN signaling, we investigated both upstream and downstream components of the pathway. Engagement of the IFNAR receptors with IFN-β stimulates the phosphorylation of STAT1 and STAT2. Phosphorylated STAT1 and STAT2 form a complex with IRF9 to translocate into the nucleus and initiate the expression of ISGs such as USP18.^31^ Examination of the transcription factors STAT1 and STAT2 phosphorylation status (Figure 3C), and the expression level of USP18 (Figure 3D, E), revealed no alterations in response to USP24 depletion, suggesting that USP24 does not interfere with upstream or downstream Type I IFN signaling. Furthermore, to assess whether the enzymatic activity of USP18 is altered in USP24-depleted cells, we labeled the cell extracts from USP24-depleted and control cells treated with IFN-β with Rho-hISG15_CTD_-PA probe, followed by fluorescence scanning and immunoblot analysis. Depletion of USP24 also did not impact the activity of USP18 (Supplementary Fig. 4). Overall, these findings indicated that depletion of USP24 enhances IFN-β induced ISGylation by affecting Type I IFN production/secretion and thereby amplifying IFNAR signaling. Importantly, the loss of USP24 cannot be fully compensated by USP18 activity, highlighting the specific role of USP24 as a de-ISGylase even in the presence of USP18.

### Quantitative proteomics identifies discrete substrates of USP24 for deISGylation

To identify relevant ISG15-modified substrates specifically targeted by USP24, we performed a proteome-wide analysis in HeLa cells with and without USP24 depletion under non-stimulated and IFN-β stimulated conditions. Considering USP24’s canonical function as deubiquitinase, we devised a workflow to identify ISGylated proteins targeted by USP24 specifically (Figure 4A). To achieve this, we employed RNAi-mediated depletion of USP24 and used scrambled siRNA (siC) as a comparative control, across both non-stimulated and IFN-β stimulated conditions. We conducted analyses in triplicates for four distinct scenarios: (I) Total Proteome to investigate USP24-dependent changes in protein levels, (II) enrichment with the di-Gly antibody recognizing Lys-ε-Gly-Gly remnants of proteins modified by Ub or Ubls (Nedd8 and ISG15) following trypsin digestion (GG-peptidome), (III) enrichment with ISG15-specific antibody (ISG15 interactome) and (IV) enrichment with the di-Gly antibody (GG-peptidome) after USP2 deubiquitinase treatment to exclude all ubiquitinated peptides (Refined ISGylome) (Figure 4A). The efficiency of sample preparation was evaluated by immunoblot analysis against indicated antibodies (Supplementary Fig. 5). A total of 7,610 unique proteins in the total proteome analysis (Supplementary Data S1), 8,169 unique Lys-ε-Gly-Gly remnant peptides (Supplementary Data S2), and 4,410 unique proteins enriched in the ISG15 interactome (Supplementary Data S3) were identified. To elucidate global similarities and differences among replicate samples and detect hidden patterns across various conditions, we employed principal component analysis (PCA) on the total proteome, GG-peptidome, and ISG15 interactome, as well as hierarchical cluster analysis on the total proteome. The PCA profile revealed robust clustering of replicates from different treatments (Supplementary Fig. 6A-C). Furthermore, hierarchical clustering analysis showed different protein expression profiles across various conditions (Supplementary Fig. 6D), including IFN-β protein patterns in siUSP24 versus siC controls (Supplementary Fig. 7A, B). Cross-comparative analysis of the USP24-dependent ISG15 interactome and GG-peptidome revealed multiple unique peptides from eleven proteins that were specifically enriched in both the ISG15 interactome and GG-peptidome (but not upon USP2 treatment) (green box in Figure 4B, C). These included Moloney Leukemia Virus 10 Protein (MOV10), Interferon Induced Protein With Tetratricopeptide Repeats 2 (IFIT2), Tryptophanyl-TRNA Synthetase 1 (WARS1), SH3 Domain-Containing Protein 19 (SH3D19), Erythrocyte Membrane Protein P55 (MPP1), UV Radiation Resistance-Associated Gene Protein (UVRAG), Cystathionine Gamma-Lyase (CTH), Bridging integrator 3 (BIN3), Interferon-Induced GTP-Binding Protein Mx2 (MX2), E3 Ubiquitin-Protein Transferase MAEA (MAEA) and Ribonuclease H2 subunit A (RNASEH2A). These proteins exhibited a ≥ 1.5 fold difference of the Log_2_ intensities in both datasets in comparison of siUSP24 treated with IFN-β versus siC treated with IFN-β (Figure 4B). Additionally, cross-comparison of the GG-peptidome analysis between IFN-β stimulated siUSP24 samples treated with USP2 versus untreated, and the ISG15 interactome analysis between IFN-β stimulated siUSP24 versus IFN-β stimulated siC also identified that the same proteins showed a ≥ 1.5 fold difference of the Log_2_ intensities for the ISG15 interactome but not for USP2-treated GG-peptidome (Figure 4C). Volcano plots of the ISG15 interactome and GG-peptidome data showed that only 7 out of the 11 proteins (MOV10, WARS1, SH3D19, UVRAG, CTH, BIN3, and MX2) were significantly enriched in siUSP24 samples treated with IFN-β compared to siC samples treated with IFN-β (Supplementary Fig. 7C-F), suggesting these seven proteins as high-confidence candidates of USP24-dependent deISGylated hits. Notably, MOV10 showed significant enrichment in siUSP24 samples compared to siC samples without treatment of IFN-β, implying that MOV10 may be a substrate of USP24 deISGlyase and possibly deubiquitinase activity (Supplementary Fig. 7D, F).

**Figure 4:**
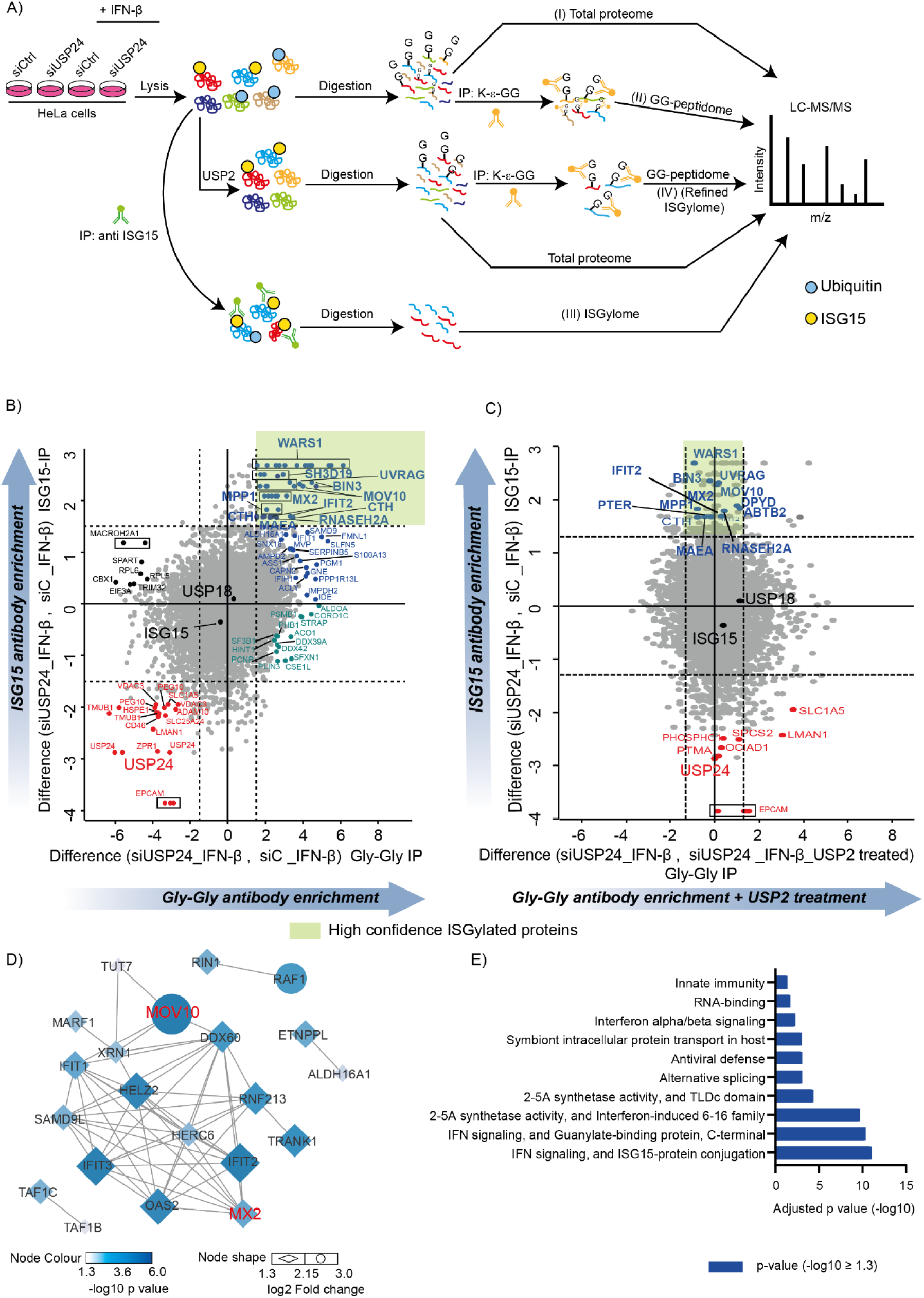
Quantitative proteomics identifies specific substrates of USP24 for deISGylation. **(A)** Schematic representation of the overall quantitative proteomic strategy deployed to investigate the USP24-dependent ISGylome and to identify specific substrates targeted by USP24 for deISGylation. Controls taken for the experiment: -/+ IFN-β stimulation for 48 h and with/without USP24 knockdown in HeLa cells (Biological replicates, n=3 for all experiments). *Overview:* Total proteome analysis (I), GG-peptidome by GG-modified peptides enrichment with di-Gly motif antibodies (II), ISG15 interactome followed endogenous ISG15 immunoprecipitation (II), ISGylome after recombinant USP2 treatment followed by di-Gly immunoprecipitation (IV). **(B)** Scatter plots of the cross-comparative analysis of ISG15 immunoprecipitation (IP) vs. di-Gly IP. Proteins indicated in the green box are significantly enriched by the ISG15 and di-Gly antibodies. Dashed lines indicate log2 ≥ 1.5, n=3 independent experiments. **(C)** Scatter plots of the cross-comparative analysis of di-Gly immunoprecipitation (siUSP24, +IFN-β, no USP2 treatment versus siUSP24, +IFNβ, with USP2 treatment) versus ISG15 immunoprecipitation (siUSP24, +IFN-β versus siC, +IFN-β). Dashed lines indicate log2 ≥ 1.5, n=3 independent experiments. The proteins shown in the green box are significantly enriched by the ISG15 antibody in siUSP24 samples compared to siC samples but not by the di-Gly antibody in siUSP24 samples treated with recombinant USP2 compared to untreated siUSP24 samples. See also Supplementary Fig. 6 and 7. **(D)** Network analysis of proteins significantly enriched by the ISG15 antibody (ISGylome). Networks were generated using the STRING database of protein-protein interactions (confidence cutoff = 0.4, default), available as a plug-in for Cytoscape 3.9.1. Node color has been set as per continuous mapping of significant enrichment as per p-value (-log10 ≥ 1.3), and the node shape (diamond/oval) has been adjusted based on fold change (log_2_ ≥ 1.3). Two of the high confidence hits targeted by USP24 for deISGylation are highlighted in red. **(E)** GO enrichment analysis of the USP24-dependent ISGylome. The bar graph shows the most significantly overrepresented GO terms, highlighting the involvement of the analyzed proteins in various biological processes analyzed using the annotated human proteome.

To gain a global insight into the ISGylome of USP24 and the associated cellular functions, we performed a network analysis of the hits enriched by the ISG15 specific antibody (ISG15 interactome), using Cytoscape 3.9.1 with STRING app (Confidence cutoff: 0.4, default). We selected hits (siUSP24, IFN-β treated – siC, IFN-β treated) based upon their p-value (-log_10_ ≥ 1.3) that represents proteins significantly enriched by the ISG15 antibody. Furthermore, we filtered based on fold change (log2 ≥ 1.3) that showed 56 proteins, which are at least 2.5 times more enriched by the ISG15 antibody upon USP24 depletion (Supplementary Data S3, S4). These 56 proteins were analyzed on Cytoscape, among which 21 proteins were found to be interconnected. This included MOV10 and MX2, two of the hits identified as potential substrates of USP24-mediated deISGylase activity (Figure 4D). To explore the biological importance of the networked proteins, we subsequently performed a functional enrichment analysis of the network, where ISG15 conjugation and IFN-I signaling came up to be the most significant. Overall, our network analysis suggests that the USP24-dependent ISGylome is associated with IFN signaling and ISGylation mechanism, where MOV10 was selected as an interesting target of USP24 deISGylation on the basis of its higher p-value and fold change compared to MX2 in the network analysis (Figure 4D, E).

### USP24 negatively regulates innate immune response by deISGylating MOV10

MOV10 is an RNA helicase involved in the regulation of gene expression by controlling mRNA decay pathways.^32^ It is also involved in the regulation of IFN signaling and antiviral responses against RNA virus^25^, but the underlying mechanism of its regulation is unknown. After observing that MOV10 is notably enriched as an ISGylated substrate in a USP24-dependent manner, we further investigated to ascertain whether MOV10 serves as a direct and specific target of USP24 deISGylase function. To this end, a hit validation experiment was carried out in USP24-depleted (siUSP24), USP24KO, and WT (siC) HeLa cells, both under IFN-β stimulation or basal conditions. To discriminate between ISG15 and Ub modifications on MOV10 and USP24 specificity towards MOV10 as deISGylase, USP18- and ISG15-depleted HeLa cells were also included as controls. The analysis showed a noticeable upward shift in the MOV10 band in USP24-depleted and USP24KO cells compared to control cells, particularly evident under IFN-β stimulation, indicating USP24-dependent modification of MOV10 under such conditions (Figure 5A). Conversely, depletion of ISG15 or USP18 did not induce any visible shift in the MOV10 band relative to control HeLa cells, thus confirming USP24-dependent ISGylation of MOV10 under IFN stimulation. To further prove the direct deISGylase activity of USP24 on MOV10, we performed a biochemical characterization of ISGylation on MOV10 by reconstituting the expression of either USP24WT or USP24 catalytically inactive Cys1698Ala mutant (USP24CA) in USP24KO HeLa cells. Ectopic expression of USP24WT but not of USP24CA mutant significantly reduced ISGylated MOV10 levels in USP24KO HeLa cells (Figure 5B, C), confirming that catalytical activity of USP24 as de-ISGylase determines the cleavage of ISG15 from MOV10.

**Figure 5:**
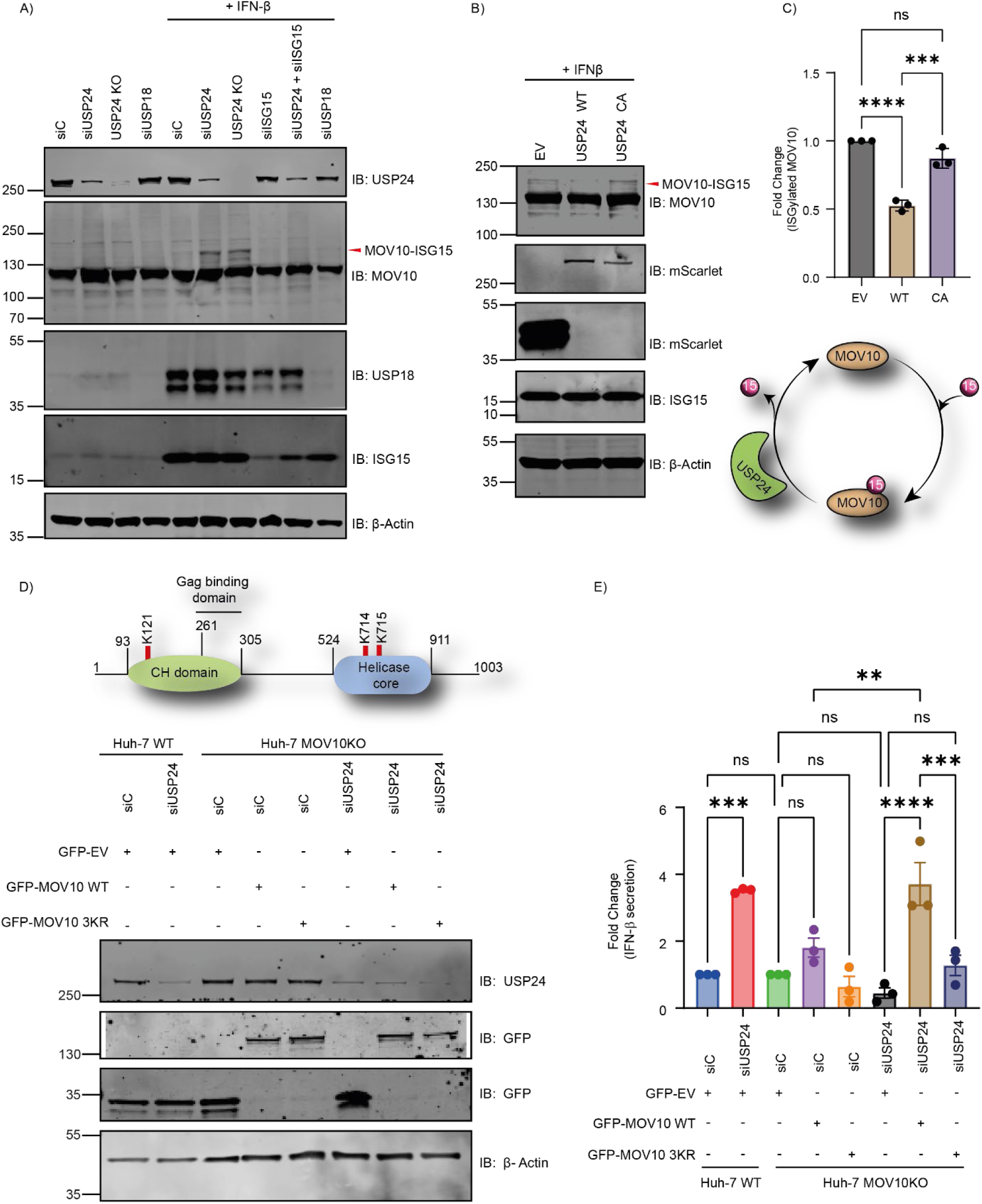
USP24 de-ISGylates MOV10 to negatively regulate IFN-I production/secretion. **(A)** Validation of MOV10 as a specific target of USP24 for de-ISGylation. WT HeLa cells transfected with the indicated siRNA oligos and USP24KO HeLa cells treated with or without IFN-β. ISGylation of MOV10 was assessed by Immunoblot against MOV10 and is observed as an upward shifted band of MOV10, corresponding to MOV10-ISG15 conjugate. Immunoblots of USP24, USP18 and ISG15 show appropriate depletion of these proteins. β-Actin serves as loading control. **(B)** ISGylation status of endogenous MOV10 as a function of USP24 de-ISGylase activity. HeLa cells expressing USP24WT, catalytically inactive mutant USP24CA, or Scarlette empty vector (EV) were assessed by immunoblot against MOV10, ISG15, and Scarlette. β-Actin is used as loading control. **(C)** Quantification of ISGylated MOV10 in the presence of EV, USP24WT, or USP24CA. Fold change was calculated by normalizing the data to EV, n=3 independent experiment. A summary of B and C is schematically illustrated at the bottom of C. **(D)** Schematic representation of the MOV10 structure showing mutated lysine residues (K121, K714, K715) located on the CH domain and the Helicase core domain at the top. See also Supplementary Fig. 8. Immunoblot analysis of depletion of USP24 and reconstitution of GFP-tagged MOV10 WT and MOV10 3KR in both WT and MOV10 KO Huh7 cells. β-Actin is used as a loading control. **(E)** Graph showing the fold change in IFN-β secretion based on the abundance of USP24 and MOV10WT/3KR in the presence or absence of USP24. IFN-β secretion in the cell culture media was determined using human IFN-β specific ELISA through a bioluminescence readout. Bar graphs report the mean, and error bars reflect ± SEM. Samples are normalized as per the siC of respective cell lines. All significant values were calculated using One way ANOVA: **p < 0.05, ***p < 0.001, ****p < 0.0001, ns = not significant.

MOV10 was previously shown to enhance IFN-β secretion^25^ whereas USP24 was reported as a negative regulator of IFN production/secretion.^26^ We hypothesized that the reversible ISGylation of MOV10, regulated by USP24 deISGylase activity, may govern IFN production and secretion. To address this, we revisited our GG-peptidomics data and identified two unique MOV10-derived peptides that were only enriched in USP24-depleted cells after IFN-β and not processed by USP2. These peptides contain two lysine residues with Gly-Gly signatures (Supplementary Fig. 8), indicating them as the bonafide ISGylation sites of MOV10 targeted by USP24. Based on this information, we generated a construct expressing GFP-tagged ISGylation-deficient MOV10 mutant by mutating lysine residues K121 and K714 to arginine. To avoid the potential hopping of ISGylation, we additionally mutated K715 due to its close proximity to K714, henceforth referred to as the MOV10 3KR mutant. Subsequently, we used the enzyme-linked immunosorbent assay (ELISA) to measure the IFN-β secretion from the Huh7 WT and MOV10KO cells transfected with siUSP24 or siC following 24 h of poly(I:C), a synthetic analog of double-stranded RNA (dsRNA), treatment. Depletion of USP24 significantly increased IFN-β secretion in Huh-7 WT cells but not in the Huh7 MOV10KO cells (Figure 5D, E), indicating a regulatory connection between USP24 and MOV10 in IFN-β secretion. To understand whether this regulation of IFN-β secretion depends on the reversible ISGylation of MOV10, we reconstituted the expression of MOV10 WT and MOV10 3KR mutant in MOV10KO Huh7 cells. Interestingly, ectopic expression of MOV10 WT significantly increased IFN-β secretion in USP24-depleted MOV10KO cells, while a slight increase was observed in control cells (siC). However, the ISGylation-deficient mutant, MOV10 3KR, was unable to rescue the effect of loss of endogenous MOV10 on IFN-β secretion in both USP24 depleted and control cells. Overall, these findings shed light on the intricate interplay between USP24, MOV10 ISGylation, and IFN-β secretion, highlighting a novel mechanism regulated by USP24 de-ISGylase activity towards MOV10 activation (Figure 6).

**Figure 6.**
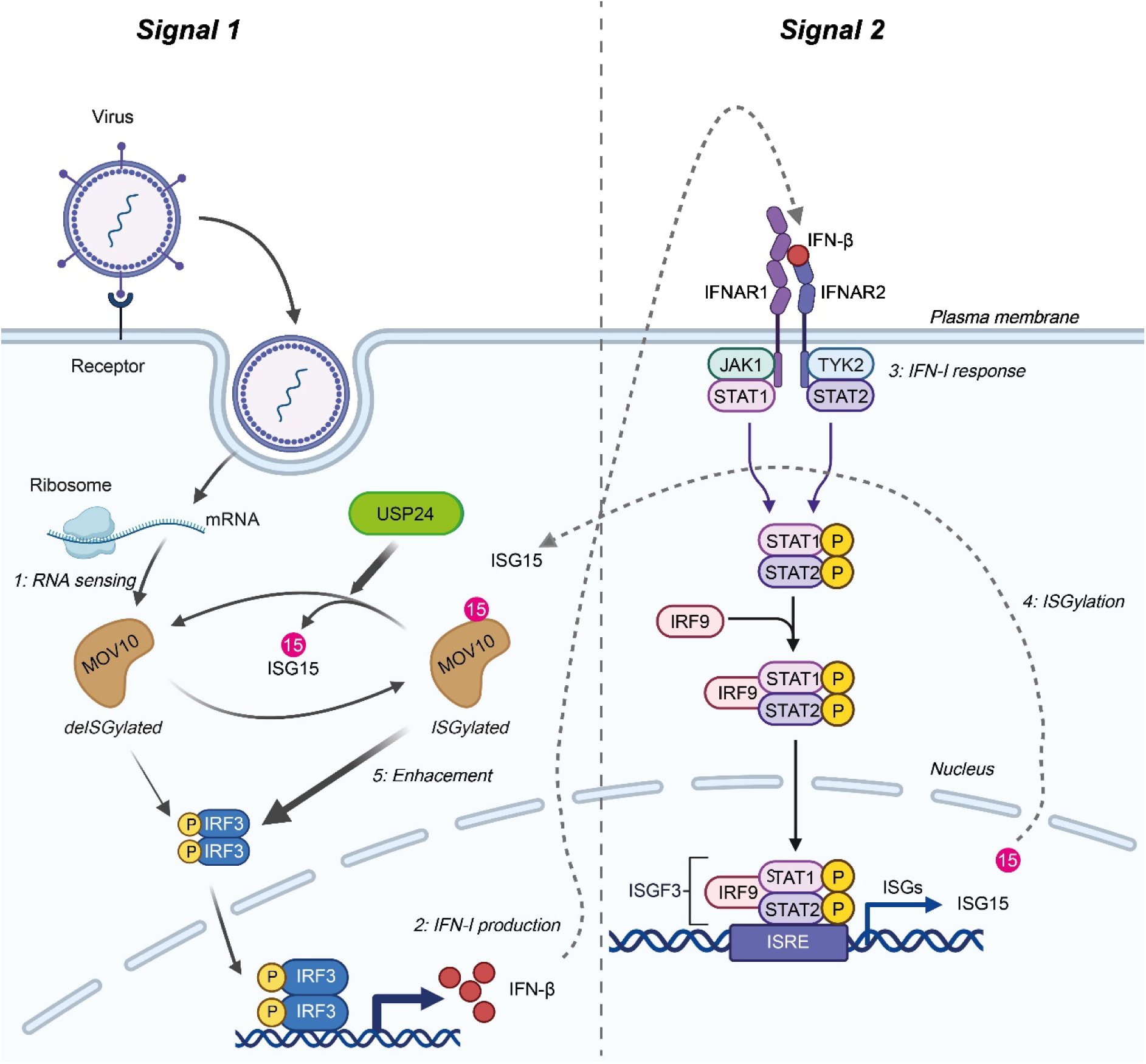
Proposed model for the role of USP24 dependent reversible ISGylation of MOV10 in IFN-β secretion and innate immune response. Viral RNA/poly(I:C) induces IFN-I production and, at a later stage, ISG15 expression in cells. ISGylation of MOV10 is necessary for it to augment IFN-β secretion by modulating IRF3 signal, acting as a positive feedback loop. In the presence of USP24, MOV10 gets de-ISGylated, resulting in reduced IFN-β secretion. On the contrary, in the absence of USP24, ISG15 conjugation on MOV10 remains, which enhances IFN-β secretion levels in Signal 1. This IFN-β is secreted and further interacts with the IFNAR receptors, generating a STAT-dependent downstream signal (Signal 2) and producing more Interferon Stimulated Genes (ISGs), including ISG15, thereby alleviating the innate immune response to the viral RNA. Adapted from “ASC-09: Potential Repurposed Drug Candidate for COVID-19” and “Interferon pathway” templates by BioRender.com (2024). Retrieved from https://app.biorender.com/biorender-templates.

## DISCUSSION

DUBs are crucial regulators of various cellular processes, including the innate immune response.^33, 34^ Typically, DUBs specifically target Ub. However, with the advancement of activity-based tools using Ub and Ubls, instances of cross-reactivity with other Ub-like proteins have been reported.^21,35^ A first glimpse of the existence of DUB cross-reactivity with ISG15, a Ub-like post-translational modifier and an important signaling protein in immunity-related pathways,^36^ was identified in an activity-based protein profiling screen involving 22 recombinant human DUBs from the USP protease family.^17^ Subsequently, USP21 was reported to process both poly-Ub and ISG15.^18^ Recent studies have identified USP16 and USP36 as ISG15 cross-reactive DUBs. ^20,21^ Additionally, our research showed that USP16 de-ISGylase activity targets ISGylated proteins involved in (immuno)metabolism. ^19^ In the current study, we expanded this panel of ISG15 cross-reactive DUBs by identifying USP24 through a pulldown experiment using our Biotin-ISG15-PA probe followed by LC-MS/MS analysis. USP24, a member of the cysteine protease USP DUB family, is gaining attention due to its broad biological functions related to its deubiquitinating activity.^37^ However, its deISGylation activity has not been previously explored. USP18, which is an ISG itself, is the primary protease for ISG15 and can process nearly all ISGylated substrates. This raises the question about the necessity of ISG15 cross-reactive DUBs in the presence or absence of USP18: Are there ISGylated substrates that USP18 does not target? If so, what is the biological significance of this cross-reactivity? To address these questions, we investigated the deISGylase activity of USP24, identifying specific substrates targeted by USP24 for de-ISGylation. Our detailed exploration reveals a novel function of USP24 as a deISGylase in the innate immune response, highlighting a new mechanism of immune regulation by cross-reactive DUBs.

The ISG15 cascade initiates from the processing of pro-ISG15 to mature ISG15, which is essential for ISG15 conjugation to substrates. Although USP18 is the primary enzyme responsible for processing pro-ISG15, its depletion does not impede the maturation of pro-ISG15 to ISG15 in cells, suggesting the involvement of other enzymes.^20,38^ As previously reported for USP16,^20^ we found that recombinant USP24 can also process pro-ISG15 into mature ISG15. This highlights the role of cross-reactive DUBs, which may compensate for the loss of USP18 or enhance the process of pro-ISG15 processing to mature ISG15. We further investigated the catalytic potential of USP24 towards ISG15 conjugated substrates using several *in vitro* ISG15_CTD_ and ubiquitin-based substrates. Full-length USP24 actively processed both ubiquitinated and ISGylated substrates. Intriguingly, when the catalytic domain of USP24 was isolated, its activity towards ISG15 was abrogated while retaining its activity towards ubiquitinated substrates. This potentially indicates the presence of other domains on USP24 to be required for ISG15 recognition by the catalytic domain or possibly the catalytic domain exhibits such minimal activity on ISG15_CTD_ substrates that it is barely detectable in *in vitro* assays.

To gain a comprehensive understanding of USP24 and ISG15 biology, the investigation was extended to mammalian cells. Depletion of USP24 in HeLa cells led to significant accumulation of ISG15 conjugates upon IFN-β stimulation. This observation was consistent in both USP24 knockout HeLa cells and siRNA-mediated USP24 depletion in HAP-1 cells, indicating a catalytic relationship between USP24 and ISG15-conjugated proteins. Further analysis confirmed that, unlike USP18,^39^ USP24 is neither an ISG nor does it affect Type-I Interferon signaling or USP18 itself, indicating that USP24 has a specific and independent role as a deISGylase. This encouraged us to explore the cellular substrates deISGylated by USP24 and investigate whether they overlap with the substrates targeted by USP18. However, as USP24 primarily functions as deubiquitinase, distinguishing ISGylated substrates from ubiquitinated ones specifically targeted by USP24 posed a challenge. We addressed this issue by performing a proteomic-wide analysis in the absence and presence of USP24 under non-stimulated and IFN-stimulated Hela cells, where we conducted four different workflows, including total proteome, the di-Gly antibody enrichment of proteins modified by Ub, Nedd8 or ISG15 following trypsin digestion, enrichment with the ISG15-specific antibody (ISG interactome) and including an additional step of treating the samples with recombinant USP2 followed by di-Gly antibody enrichment, which eliminated most of the ubiquitinated substrates and specifically revealed ISGylated peptides affected by USP24 depletion.^40^ Cross-comparisons of these proteomic analyses revealed seven ISGylated substrates potentially targeted by USP24, which include MOV10, WARS1, SH3D19, UVRAG, CTH, BIN3, and MX2. Bioinformatic analysis and subsequent in-cell validation confirmed MOV10 as a USP24-specific substrate targeted for deISGylation, even in the presence of USP18. This indicates the existence of ISGylated substrates that are not processed by USP18 but require other cross-reactive DUBs like USP24. A Gene Ontology (GO) analysis of the USP24-dependent ISGylome emphasized IFN-I signaling, raising the question about the underlying biological importance of the USP24-MOV10-ISG15 axis in innate immunity.

MOV10 is an RNA helicase known for its role in suppressing viral replication.^24^ Previous studies have identified a direct correlation between MOV10 and IFN-α in patients with Chronic Hepatitis B virus (HBV) infection. HBV-infected patients exhibited significantly lower mRNA levels of both MOV10 and IFN-α compared to a control group.^41^ IFN-β, another IFN-I, production and secretion is a critical event in the innate immune response, playing a pivotal role in activating immune cells and stimulating adaptive immunity.^42^ Another study highlighted the influence of MOV10 on IFN pathways by demonstrating that MOV10 enhances IFN-β production, thereby accelerating cell antiviral activity.^25^ The study demonstrated that in the presence of MOV10, upon viral infection, there is increased nuclear localization of the transcription factor interferon regulatory factor 3 (IRF3), which mediates IFN-β mediated immune response.^25^ However, the mechanism by which MOV10 is regulated to modulate IFN-β secretion was not understood. Our investigation suggests that ISG15 modification of MOV10 can modulate IFN-β secretion following poly (I:C) treatment in cells mimicking RNA virus infection. USP24 deISGylates MOV10, reducing MOV10-mediated acceleration of IFN-β secretion and acting as a negative regulator of the IFN response. This suggests that inhibiting USP24 could prevent the deISGylation of MOV10, thereby elevating cellular IFN-β production/secretion levels (signal 1) and indirectly enhancing type I IFN receptor signaling (signal 2) (Figure 6).

Additionally, IFN-β has previously been explored therapeutically. A study identified several small molecules that significantly inhibited SARS-CoV2 replication *in vitro* by modulating the IFN-β induced host signatures, though the underlying molecular mechanism has not been reported yet.^43^ Moreover, other studies have identified IFN-α and IFN-β as anticancer immunotherapeutics that improve the outcome of diverse malignancies by influencing the immune, vascular, and apoptotic processes through transcription of pleiotropic factors. Thus, new insights into IFN regulation offer opportunities to combine interferon activation with other therapies in cancer treatment.^44^ Therefore, USP24 inhibitors have the potential to serve as adjunct antiviral therapy and boosters of cancer immunotherapy. This finding encourages further research to screen for USP24 inhibitors^45^ and to identify other cross-reactive DUBs, exploring their biological roles. Such studies could open new avenues for understanding the role of DUBs in crucial pathways involved in various diseases and identify novel drug targets. Our extensive proteomic data also highlighted other Ub/ISG15-conjugated targets of USP24 besides MOV10. Investigating their biological functions could reveal further important roles of USP24, not only in innate immunity but also in other cellular mechanisms.

## MATERIALS AND METHODS

### Cell Culture

EL4 cells were cultured in Dulbecco’s modified Eagle’s medium (DMEM) (Gibco) supplemented with 2mM Glutamine and 10% fetal bovine serum (FBS). HEK293T (Cat# ATCC® CRL-3216™), HeLa, Huh-7 WT, and Huh-7 MOV10KO cells were cultured in Dulbecco’s modified Eagle’s medium (DMEM) (Gibco), and HAP1 cells were cultured in Iscove’s modified Dulbecco’s medium (IMDM) (Gibco). Both types of medium were supplemented with 8% FCS. Cells were grown at 37 °C in a humidified 5% CO_2_ incubator. All cell lines have been authenticated and routinely tested for mycoplasma.

### DNA constructs

A full-length USP24 plasmid was provided by Dr. Feng Gong (University of Miami Health System).^46^ It was recloned into an in-house-available mScarlet-C1 vector with a CMV promoter. Subsequent clones were created in different vectors according to the expression system following the *In Vivo* Assembly (IVA) cloning technique.^47^ MOV10 plasmid was purchased from Addgene (# 10976) and recloned into an in-house mGFP-C1 vector following the IVA technique. For site-directed mutagenesis, a mixture of template DNA, 1x Pfu buffer, 10 mM dNTPs, 125 ng forward and reverse primers with the desired mutation(s), 1 µL of TURBO Pfu Polymerase (Agilent), and 50 ng template DNA was combined with autoclaved MilliQ water to a final volume of 50 µL. PCR was done using the following program: 98 °C for 3 mins, 60 °C (depending on primer Tm) for 20 sec, 72 °C for 30 sec/kb of template size, 72 °C for 5 mins, 4 °C on infinite hold. 40 µL of the amplified PCR product was incubated with 1 µL DpnI (Thermo Fischer Scientific) for 2 h at 37 °C and transformed into competent DH5α bacterial strain. All modified sequences were verified by sequencing. Primers used are enlisted in Supplementary Table 1 and were purchased from Sigma-Aldrich.

### Transfection

For DNA transfection, HEK293T and HeLa cells were seeded to achieve around 60% confluency the following day, while Huh7 cells were reversibly transfected as described previously.^48^ DNA was transfected using PEI (polyethylenimine, Polysciences Inc., Cat# 23966). The transfection mixture included 200 µL DMEM without supplements, the required amount of plasmid of interest, and PEI (1 mg/mL). DNA: PEI ratio was always maintained at 1 : 3 (1 µg Plasmid : 3 µg PEI). The mixture was incubated at room temperature for 20 minutes without shaking and then added to the cells dropwise. Cells were monitored after 24 h and then collected.

For siRNA transfection, human USP24 (On-TARGETplus Cat# J-006073-11), ISG15 (OnTarget Plus # L-004235-03-0005) and MOV10 (OnTarget Plus # L-014162-00-0005)specific siRNAs were purchased from Dharmacon, while human USP18 specific siRNA (#AM16708) was purchased from ThermoFisher Scientific. Silencing was performed in HeLa and HAP-1 cells. In 24-well format, 50 μL of siRNA (500 nM stock) was incubated with 1 μL Dharmafect reagent 1 (Dharmacon) diluted in 49 μL medium without supplements (total volume of 100 μL transfection mix). Incubation was done at room temperature with gentle shaking for 20 mins. A total of 2 x 10^4^ cells were resuspended in 400 μL of growth medium without antibiotics and added to the transfection mix. The final volume, along with the transfection mix, was 500 μL per well. 24 h post-transfection, cells were treated with IFN-β (1:1000 or 1000 U/mL) and incubated for 48 h. Cells were cultured for a total of 3 days prior to further analysis.

### CRISPR/Cas9- mediated knockout of USP24

gRNA sequences targeting the USP24 gene (gRNA1: GCTCGTAGCCGCCGTAGTCG, gRNA2: GAACGACATTAACGAGGCCG, gRNA3: CTACGGCGGCTACGAGCCCA, gRNA4: ACTGCTCACCAACGAGCGGC) were purchased from Integrated DNA Technologies (IDT) and cloned into the PX459 plasmid (containing the Cas9 gene and a puromycin resistance gene) using IVA cloning^47^. This plasmid was transiently transfected into HeLa cells using Effectene and the following day, cells were selected with 200 µg/mL puromycin for 72 h. Cells were then diluted and maintained in a 15 cm dish, allowing colonies to grow distinctly. Isolated colonies were collected, expanded, and analyzed for the loss of USP24 by Western blot. Huh-7 MOV10 CRISPR knockout cells were obtained from Dr. Sonia Zuñiga.^48^

### SDS-PAGE and Immunoblotting

Samples were resolved on precast Bis-Tris NuPAGE gels (Invitrogen, 4-12%, 12%) and Tris-Acetate gels (Invitrogen, 3-8%). MOPS buffer (Invitrogen Life Technologies, Carlsbad, USA) for Bis-Tris gels and Tris-Acetate buffer (Invitrogen Life Technologies, Carlsbad, USA) for Tris-Acetate gels was used as running buffer. Proteins were transferred to a nitrocellulose membrane (Protan BA85, 0.45 μm, GE Healthcare) at 300 mA for 2.5 h. The membranes were blocked in 5% milk (skim milk powder, LP0031, Oxiod) in 1× PBS (P1379, Sigma-Aldrich), incubated with a primary antibody diluted in 5% milk in 0.1% PBS-Tween 20 (PBST) overnight, washed three times for 10 min in 0.1% PBST, incubated with the secondary antibody diluted in 5% milk in 0.1% PBST for 30-45 min, and washed three more times in 0.1% PBST. In the case of pSTAT1 and pSTAT2 blots, milk was replaced with BSA, and the buffers used were TBS and TBS-T. Immunoblot signals were visualized with a LI-COR Odyssey Fx laser scanning fluorescence imager and analyzed using LI-COR Image Studio Lite software.

### Antibodies

Primary antibodies used for the study were: rabbit anti-USP24 (Abcam, #ab129064), rabbit anti-USP18 (D4E7) (Cell Signalling Technology, #4813), rabbit anti-ISG15 (Invitrogen, #50546), rabbit anti-MOV10 (Proteintech #10370-1-AP), rabbit anti-mGFP^49^, rabbit anti-mRFP^49^, mouse anti-β Actin (Sigma-Aldrich, #A5441). STAT1 (D4Y6Z) (Cell Signalling Technology, #14995), phosphoSTAT1 (Y701) (58D6) (Cell Signalling Technology #9167), STAT2 (D9J7L) (Cell Signalling Technology #72604), phosphoSTAT2 (Y690) (D3P2P) (Cell Signalling Technology #88410). Secondary antibodies used were: IRDye 800CW goat anti-rabbit IgG (H + L) (Li-COR, Cat# 926-32211, 1:5000), IRDye 800CW goat anti-mouse IgG (H + L) (Li-COR, Cat# 926-32210, 1:5000), IRDye 680LT goat anti-rabbit IgG (H + L) (Li-COR, Cat# 926-68021, 1:20,000), and IRDye 680LT goat anti-mouse IgG (H + L) (Li-COR, Cat# 926-68020, 1:20,000).

### Stimulation of cells

Lyophilized recombinant human IFN-β (PeproTech # 300-02BC-B) was diluted into 200 µL of sterile MilliQ water or LAL water to make the stock. 10 µL aliquots were prepared and stored at −80 °C. All treatments were done from this stock. Cells were treated with 1000 U/mL of recombinant human IFN-β stock for the indicated time period.

Viral dsRNA-induced infection in cells was mimicked by treating the cells with 2 µg/mL of poly(I:C) LMW (InvivoGen, Cat. Code: tlrl-picw) dissolved in sterile PBS. Poly(I:C) (at a final concentration of 2 µg/mL) was mixed with PEI (1:3) following incubation at room temperature for 20 mins. The mixture was added to the cells and left for the required time as per the experiment.

### ISG15 conjugate detection

Formation of ISG15 conjugates was detected using ISG15 specific antibody (Invitrogen, ISG15 Recombinant Rabbit Monoclonal Antibody (7H29L24), Catalog # 703131) after depletion of USP24 followed by IFN-β treatment. Cells were collected after 48h following IFN-β treatment. Cells were lysed using Lysis buffer (50 mM Tris-HCl pH 7.4, 150 mM NaCl, 0.5% TRITON X-100, 1 tablet of complete EDTA-free protease inhibitor cocktail (Roche) per 10 mL buffer). Protein quantification was done using BCA assay (Thermo Scientific™, Pierce™ BCA Protein Assay Kits, Catalog number: 23225) performed following manufacturer’s protocol and an equal amount of protein was loaded and run on SDS-PAGE gel (Invitrogen™, NuPAGE™ Bis-Tris Mini Protein Gels, 4–12%, 1.5 mm, Catalog number: NP0336BOX). Proteins were transferred to the Nitrocellulose membrane, and immunoblotting was performed as described. ISG15 conjugates were quantified using LI-COR Image Studio Lite software. The intensity of ISG15 conjugated bands was normalized to the β-actin bands, and fold change between siC and siUSP24 treated samples in ISG15 conjugation was determined.

### Activity-based probe labelling and pull-down assays

Biotin-tagged full-length mouse ISG15-PA,^13^ untagged human full-length ISG15-PA, Rhodamine-tagged C-terminal domain ISG15_CTD_-PA, Biotin-tagged human C-terminal domain ISG15_CTD_-PA, and Rhodamine-tagged Ub-PA probes were used^50^. For DUB labelling with probes in cell lysates, cell pellets were resuspended in Lysis buffer (50 mM Tris-HCl pH 7.4, 150 mM NaCl, 2 mM TCEP, 0.5% TRITON X-100, 1 tablet of complete EDTA-free protease inhibitor cocktail (Roche) per 10 mL buffer). Lysis was done by sonication using a Bioruptor sonicator (Diagenode) at medium intensity for 3 bursts with an ON/OFF cycle of 2 seconds at 4 °C. Cell lysate was clarified by centrifugation at 4 °C for 20 mins at a speed of >12000xg. 20-30 μg of clarified cell lysate in 20 μL was treated with the indicated probe (final probe concentration 1 μM) at 37 °C for 30 min. Reactions were stopped by adding LDS (lithium dodecyl sulfate) sample buffer (Invitrogen Life Technologies, Carlsbad, CA, USA) containing 2.5% v/v β-mercaptoethanol and boiling for 7 minutes. Samples were loaded and run on SDS-PAGE gel as described in the “SDS-PAGE Electrophoresis and Immunoblottings” section. Enzymes labeled with fluorescent probes were visualized by in-gel fluorescence scanning using Typhoon FLA 9500 imaging system (GE Healthcare Life Sciences), where indicated. The Rhodamine channel (ex: 570nm, em: 590nm) was used to detect the probes, and the Cy5 channel (ex: 651nm, em: 670nm) was used to detect the protein marker (ThermoFisher Scientific PageRuler Plus Prestained protein ladder: 10-250 kDa). Following the fluorescent scan, proteins were transferred to the nitrocellulose membrane, followed by immunoblotting analysis using the indicated antibodies.

For pull-down, the medium was completely removed from the cells, followed by the addition of 300 µL lysis buffer and scraping of the cells. Cell lysis was done as described above. The supernatant was transferred to a new Eppendorf tube. 15-20 µL of supernatant was removed to blot for the input sample. The remaining supernatant was incubated with a probe (diluted in the same buffer to a final concentration of 5 µM) for 60 mins at 37 °C. Next, the total volume was completed to 1 mL with the lysis buffer. 30 µL of Neutravidin beads (Thermo Scientific™, Pierce™ NeutrAvidin™ Agarose, Catalog number: 29200) were added, followed by incubation at 4 °C on rollers overnight. Beads were collected by centrifugation at 500xg for 1 min at 4 °C and washed with wash buffer (50 mM TRIS, pH 7.5, 150 mM NaCl, 0.5% Triton X, 0.5% SDS) at least 3 times by centrifuging at 500xg for 1 min at 4 °C. Prior to the final washing, beads were transferred to a new tube to avoid background and finally centrifuged at 500xg for 3 mins at 4 °C. Beads and supernatant were stored at −80 °C until following analyses, including immunoblotting, fluorescent scanning, coomassie staining and LC/MS-MS.

For probe labeling using recombinant enzymes, the indicated concentrations of enzymes diluted in assay buffer (50 mM Tris-HCl pH 7.5, 100 mM NaCl, 0.5 mg/mL CHAPS, 5 mM TCEP/DTT) were used. Samples were incubated with a probe (diluted in the same buffer to a final concentration of 5 µM) at a 1:1 molar ratio for 60 mins at 37 °C. The reaction was quenched with LDS sample buffer, followed by boiling for 7 mins. Samples were loaded and run on SDS-PAGE gel as described in the “SDS-PAGE Electrophoresis and Immunoblottings” section. Enzymes labeled with fluorescent probes were visualized by in-gel fluorescence scanning using Typhoon FLA 9500 imaging system (GE Healthcare Life Sciences), followed by Coomassie staining.

### STAT1/STAT2 phosphorylation assay

Phosphorylation levels of STAT1 and STAT2 were monitored in wild-type and USP24 knockdown HeLa cells. Cells were treated with siRNA control and siRNA targeting USP24 (siUSP24 _Cat# J-006073-11: GGACGAGAAUUGAUAAAGA). They were allowed to grow for 72 h at 37 °C in a humidified 5% CO_2_ incubator. Following this, the cells were starved in 0.5% FCS containing DMEM media for at least 6 h. After that, the media was replaced with 0% FCS DMEM, and cells were treated with IFN-β (1:1000 or 1000U/mL) as per the depicted time points. Cells were collected in lysis buffer (see above) and subjected to BCA assay to determine total protein concentration. Sample analysis was done using SDS-PAGE and immunoblotting as described in the “SDS-PAGE Electrophoresis and Immunoblottings” section using STAT1, STAT2, pSTAT1 and pSTAT2 antibodies.

### Protein expression and purification

Recombinant His10-USP24 full-length protein was purchased from Biotechne (Catalog # E-616-050). To design the construct for USP24 catalytic domain, protein homology modeling was used using Phyre2 V2.0^51^ and SWISS-MODEL^52^ servers. This identified USP7CD (PDB: 5KYC)^53^ as a homologue of USP24CD. Based on this, the domain extensions of USP24CD were decided and the purification protocol was designed. USP24CD (aa. 1686-2062) was cloned into a pFastBac insect cell expression vector with an N-terminal 6xHis tag and a 3C protease cleavage site. The recombinant protein was expressed through the Bac-to-Bac baculovirus expression system (Invitrogen, Carlsbad, CA, USA). Cells were lysed using Lysis buffer (50 mM HEPES, pH 7.5, 500 mM NaCl, 3 mM TCEP and 1 tablet Roche Protease inhibitor per 10 mL of buffer). Lysate was clarified by centrifugation at 24,000 g for 40 mins at 4 °C. Clarified lysate was passed over Ni-NTA trap beads. Beads were washed using Wash buffer (50 mM HEPES, pH 7.5, 500 mM NaCl, 20 mM Imidazole). Protein trapped on beads were eluted using Elution buffer (50 mM HEPES, pH 7.5, 100 mM NaCl, 250 mM Imidazole, 3 mM TCEP). Samples were collected at each step of purification to monitor protein stability and yield. Selected elution were collected and dialyzed overnight in Dialyzing buffer (50 mM HEPES, pH 7.5, 100 mM NaCl, 2 mM TCEP). Dialysate was collected, concentrated, and subjected to Size Exclusion Chromatography using SEC buffer (50 mM HEPES, pH 7.5, 100 mM NaCl, 2 mM TCEP). The final product was concentrated (yielding around 320 µL of 0.2 mg/mL from 1 L of culture) and stored at −80 °C until further analyses.

### pro-ISG15 cleavage

Recombinant human pro-ISG15 protein (R&D Systems, #UL-615-500) was diluted in a buffer containing 50 mM Tris-HCl pH 7.4, 150 mM NaCl, 2 mM EDTA, 0.5 % NP40, 8 mM TCEP. 5 µM final concentration of pro-ISG15 was incubated with 0.5 µM of recombinant enzymes at 37 ^°^C over the indicated time points. The reaction was quenched by boiling with LDS sample buffer at 95 °C. Following, samples were loaded on a 12% Bis-TRIS gel (Invitrogen™, NuPAGE™ 12%, Bis-Tris, 1.0 mm, Mini Protein Gels, Catalog number: NP0341PK2) and allowed to run with MOPS buffer. Coommassie staining was done using InstantBlue® Coomassie Protein Stain (ISB1L, Abcam) to visualize the proteins.

### Fluorescence Intensity assay

Fluorogenic assay as a measure of Fluorescence Intensity (FI) was carried out using 2-fold serial dilutions of recombinant enzymes as indicated. Synthetic fluorogenic substrate (Ub-RhoMP and ISG15_CTD_-RhoMP) was used at a final concentration of 400 nM. All dilutions were made in assay buffer (50 mM Tris-HCl pH 7.5, 100 mM NaCl, 1 mM TCEP, 1 mg/mL CHAPS, 0.5 mg/mL Bovine Gamma Globulin (BGG) dissolved in MilliQ water). For the assay, 384 well microplates (Corning 3820 black, low volume, round well, flat bottom) were used. Plate reading was done on a BMG LabTech PHERAstar Plate Reader (excitation, 485 nm; emission, 520 nm) over a time period of 60 mins. The fluorescence intensity reads were plotted using Graphpad Prism software.

### Fluorescence Polarization assay

Fluorescence polarization assay was carried out using 2-fold serial dilutions of recombinant enzymes. All dilutions were made in assay buffer (50 mM Tris-HCl pH 7.5, 100 mM NaCl, 1 mM TCEP, 1 mg/mL CHAPS, 0.5 mg/mL Bovine Gamma Globulin (BGG) dissolved in MilliQ water). For the assay, 384 well microplates (Corning 3820 black, low volume, round well, flat bottom) were used. 10 µL of the enzyme dilutions were added to the wells. Synthetic substrates were dissolved in DMSO, then diluted to 400 nM stock concentration in assay buffer. The reaction was initiated by the addition of 10 µL of the indicated synthetic substrates, resulting in a 200 nM final concentration of the substrate. The fluorescence polarization was recorded every 60-90 seconds for a total of 90 mins on a BMG LabTech PHERAstar Plate Reader (excitation, 540nm; emission, 590nm) and plotted using Graphpad Prism software.

### Sample preparation for quantitative mass spectrometry analysis

HeLa cells were grown in DMEM medium supplemented with 10% FCS for label-free quantitation (LFQ) experiments. USP24 was knocked down in HeLa cells using USP24-specific siRNA, as mentioned in the "Transfection” section. Four different conditions were used: USP24 knockdown with and without Interferon-β treatment and siRNA Control with and without Interferon-β treatment. Samples were prepared in triplicate sets for each analysis and were subjected to statistical quantitation.

### Total proteome analysis

Cells were lysed using lysis buffer (50 mM Tris-HCl, pH 7.4, 0.5% NP-40, 150 mM NaCl, 20 mM MgCl_2_) containing 1 tablet of complete EDTA-free protease inhibitor cocktail (Roche) and PhosSTOP (PHOSS-RO Roche) phosphatase inhibitor per 10 mL of buffer. Protein amounts were measured using the BCA Protein assay kit (Pierce, #23225). 90 µg of protein was taken for the total proteome analysis from each sample. Samples were mixed with 2x Gel Loading buffer having 25 mM DTT. They were boiled for 10 mins at 90 °C and stored at −20 °C to be processed later. Next, samples were thawed at room temperature. A final clarification of the samples was done using S-trap mini spin columns (Protifi Innovative Omics solutions). Samples were reduced using 0.8 µL of 1.25 M DTT per 100 µL (final concentration of 10 mM DTT per sample). Alkylation of the disulfides was done using 10 µL of 200 mM Iodoacetamide (IAA) per 100 µL of sample (final concentration of 20 mM). Following this, the sample was acidified using 10 µL of 1.1% phosphoric acid (final pH of sample ≤ 1), causing the samples to change color to bright yellow. To the samples was added 620 µL of binding/wash buffer (90% Methanol, 100 mM TEAB, pH 7.55), which caused samples to regain their blue color. Samples were next loaded onto S-trap columns and centrifuged at 4000 g for 30 sec in order to allow the proteins to bind to the column. The columns were washed with 400 µL of the same buffer 4 times by centrifuging at 4000 g for 30 secs each. The residual buffer was removed by final centrifugation at 4000 g for 1 min. Finally, the filters were transferred to new 2 mL collection tubes, and 125 µL of Trypsin (1:10 wt:wt ratio) was added to the filter. Tubes were kept at room temperature without shaking for overnight digestion. Peptides were eluted from the S-trap columns using Buffer 1 (50 mM TEAB in water, pH 8.5), Buffer 2 (0.2% formic acid in water), and Buffer 3 (50% acetonitrile in water) successively. Each time, 80 µL of the buffer was applied to the column and centrifuged at 4000 g for 1 min. Eluted proteins were dried using a vacuum evaporator (SP Genevac miVac centrifugal evaporator miVacATS life sciences scientific products). Dried samples were stored at −80 °C. They were resuspended prior MS analysis in mass spectrometry grade 98% water + 2% acetonitrile + 0.1% TFA to a total volume of 15 µL.

### ISG15 interactome

Cells were lysed using lysis buffer (50 mM Tris-HCl, pH 7.4, 0.5% NP-40, 150 mM NaCl, 20 mM MgCl_2_) containing 1 tablet of complete EDTA-free protease inhibitor cocktail (Roche) and PhosSTOP (PHOSS-RO Roche) phosphatase inhibitor per 10 mL buffer. Protein amounts were measured using the BCA Protein assay kit (Pierce, #23225). 3 mg of protein was subjected to immunoprecipitation using ISG15 antibody (Recombinant rabbit monoclonal antibody: ThermoFisher scientific: 7H29L24). Lysates were incubated for 1 hr with the antibody on a rotor at 4 °C, where no antibody samples were used as negative control. Finally, 35 µL of Protein G agarose beads (ThermoFisher Scientific, Catalog# 20397) was added to the samples for immunoprecipitation. Samples were kept on the rotor overnight at 4 °C for efficient pull-down. Samples were centrifuged at 4 °C at 2000 g for 1 min. The supernatant was discarded, and beads were washed with the lysis buffer (without inhibitors) 3-4 times. The beads were eluted using 2x Gel Loading buffer with 25 mM DTT. First, elution was done using 50 µL of GLB and boiled at 90 °C for 5 mins. The next elution was done using 60 µL of GLB and boiled at 90 °C for 10 mins. Finally, both the elution sets were combined. This contains the proteins modified with ISG15. Here, samples were analyzed using a western blot to confirm the efficiency of the immunoprecipitation (Supplementary Fig. 5).

### Lys-ε-Gly-Gly antibody pull-down

Cell pellets were dissolved in 1 mL of 6 M urea diluted with 5 mL MilliQ water, reduced using dithiothreitol (4.5 mM final) for 30 min at 55 °C. This was followed by alkylation using iodoacetamide (100 mM final) for 15 min at room temperature in the dark. Protein digestion was done overnight at room temperature with trypsin (Promega V5113, 1/100 volume of 1mg/mL trypsin). 1% final concentration of TFA was added to quench the digestion. Sep-Pak C18 columns (WATERS, WAT020515) were used to purify the peptides. Purified peptides were dried using vacuum centrifugation. Peptides were later enriched using di-Gly antibody immunoprecipitation using the PTMScan Ubiquitin Remnant Motif (K-ε-GG) Kit (Cell Signaling catalog# 5562). Post immunoprecipitation, enriched peptides were desalted using Sep-Pak C18 columns (WATERS, WAT020515) and dried by vacuum centrifugation. Dried peptides were resuspended in 10 µL of mass spectrometry grade water with 2% acetonitrile and 0.1% formic acid. Resuspended peptides were stored at −20 °C until analysis.

### Refining of the ISGylome

Cells were lysed using lysis buffer (50 mM Tris-HCl, pH 7.4, 0.5% NP-40, 150 mM NaCl, 20 mM MgCl_2_, 3 mM TCEP) containing 1 tablet of complete EDTA-free protease inhibitor cocktail (Roche) and PhosSTOP (PHOSS-RO Roche) phosphatase inhibitor per 10 mL buffer. Protein amounts were measured using the BCA Protein assay kit (Pierce, #23225). Recombinant USP2 was purchased from Ubiquigent (Isoform 4: GST tagged: Catalog# 64-0014-050). 1 µM of USP2 final concentration was added to the samples and incubated for 1 h at 37 °C on a shaker at 300 rpm. For negative control, samples not treated with USP2 were taken and incubated for 1 h at 37 ^0^C on a shaker at 300 rpm.

From the final sample, 90 µg protein was used for total proteome analysis and 3 mg protein was used for Lys-ε-Gly-Gly pull-down analysis. The protocol for total proteome and K-ε-GG pull-down was the same as discussed above.

### DIA-PASEF data acquisition and DIA-NN analysis

Peptides for analysis by LC-MS/MS were loaded on Evotip Pure C18 tips (Evosep) following the manufacturer’s protocol. Briefly, tips were rinsed with 20 µL Solvent B by centrifugation, conditioned by soaking in 1-propanol, equilibrated with 20 µL Solvent A by centrifugation, loaded with 2 µL of sample and 18 µL of solvent A, then centrifuged, washed with 20 µL solvent A by centrifugation, and wetted with 100 µL solvent a and a brief centrifugation step for 10 s. All centrifugation steps were 60 s at 800 *g* unless stated otherwise.

Peptides were analysed using an Evosep One liquid chromatography system coupled to a timsTOF SCP mass spectrometer (Bruker) using the Whisper 20 samples per day method and a 75 µm x 150 mm C18 column with 1.7 µm particles and an integrated Captive Spray Emitter (IonOpticks). Data were acquired using diaPASEF^54^ methods generated by py_diAID.^55^ Total proteome and ISG15 pulldown samples were analysed using a method with 24 variable-width PASEF frames, with two ion mobility windows per PASEF frame, covering a region between 300 – 1,200 m/z and 0.6 – 1.6 V s cm^-2^. diGly samples were analysed using a method with 12 variable width PASEF frames, with two ion mobility windows per frame, covering a region between 330 – 1,100 m/z and 0.69 – 1.57 V s cm^-2^. For both methods, the accumulation and ramp times were 100 ms and the collision energy was ramped linearly over the ion mobility range: 20 – 59 eV over 0.6 – 1.6 V s cm^-2^. Raw mass spectrometry files were label-free quantified using DIA-NN (version 1.8) in library-free mode using the Uniprot proteome UP000005640 (2022) as a FASTA file.

### Bioinformatic analysis of DIA data

A peptide intensity table with the Label-free quantitation (LFQ) values was analyzed in Perseus (v1.6.2.3).^56^ Data were log_2_ transformed and filtered for identification in all three replicates in at least one group. Different peptide groups were selected for different analyses like total proteome, ISG15 IP, and diGly IP. Principal component analysis (PCA) was performed for each analysis with default settings. Intensities were *Z-scored* by subtracting the mean and used for hierarchical clustering by Euclidean distance (pre-processed with k-means, 300 clusters, 1000 iterations). Missing values were imputed from the lower end of the normal distribution (default settings). A two-sided student’s t-test with permutation-based FDR was used to calculate the significance between IFN-β treatment and control at 0.05 FDR (p-value). For a detailed protocol of data analysis in Perseus, refer to https://cox-labs.github.io/coxdocs/interactions.html.

### Gene Ontology and STRING network analysis

Cytoscape (v3.9.1)^57^ with apps STRING (protein-protein interaction, confidence cutoff = 0.4; default) was used to create and visualize the interaction networks using the UniProt gene IDs from the ISG15 IP (ISGylome) data set (siUSP24, IFN-β treated – siC, IFN-β treated) based upon their p-value (-log_10_ ≥ 1.3) and fold change (log_2_ ≥ 1.3). Gene Ontology (GO) analysis of the interconnected proteins was done using the Functional Enrichment option in Cytoscape, which generated the GO data against the human proteome. Following the GO analysis, data was exported as a “.csv” file, and the GO analysis graph was generated in GraphPad Prism5 software using the −log10 p-value (Adjusted p-value) indicated for each of the biological processes reported by the GO analysis.

### Human IFN-β bioluminescent ELISA

Cell culture supernatant was collected and clarified by centrifuging at 1000 g for 1 min. The clarified supernatant was transferred to a clean Eppendorf tube. Freshly collected media was used for better results in the ELISA as recommended in the manufacturer’s protocol. IFN-β concentration in the media was calculated through the enzyme-linked immunosorbent assay (ELISA) performed using the LumiKine™ Xpress hIFN-β 2.0 (Human IFN-β bioluminescent ELISA kit 2.0) (Cat. Code: luex-hifnbv2). The assay was performed following manufacturer’s protocol. Data is expressed as fold change of IFN-β secretion in each sample compared to the control of the respective cell line type and represents the comprehensive result obtained from at least three individual biological replicates. Calculations were done using the principles of ANOVA and represented as mean ± SEM.

### Statistical analysis

All statistical evaluations were reported on Student’s *t* test (two-tailed distribution)/ANOVA having *p < 0.05, **p < 0.01, ***p < 0.001, ****p < 0.0001 and ns: not significant. All error bars correspond to the mean ± SD or mean ± SEM (indicated in figure legends). Data were analyzed using the Microsoft Excel and GraphPad Prism5 software.

## ACKNOWLEDGEMENTS

We are grateful to Prof. Feng Gong, University of Miami Health System for providing us with the USP24 plasmid. Mass spectrometry analysis was performed in the Discovery Proteomics Facility (Target Discovery Institute) led by R. Fischer and I. Vendrell. This work was supported by the TRIM-NET project, which received funding from the European Union’s Horizon 2020 research and innovation programme under the Marie Skłodowska-Curie grant agreement No 813599. A.S was supported by ICI00026 from the NWO Gravity Program. S.D., A.P.F. and B.M.K. were supported by the Chinese Academy of Medical Sciences (CAMS) Innovation Fund for Medical Science (CIFMS), China [grant number: 2018-I2M-2-002] and by Pfizer. Work in the A.P.F. lab was funded by Ono Pharma UK Ltd. S.Z was supported by the Government of Spain (PID2022-140328OB-I00, MCIN/AEI/10.13039/501100011033/FEDER, UE). M.G received a contract from Comunidad de Madrid (PIPF-2022/SAL-GL-24549). K.P.K. received funding from the German Research Foundation (DFG), Project ID DFG 423813989/GRK2606 and under Germany’s Excellence Strategy (CIBSS–EXC-2189–Project ID 390939984).

## AUTHOR CONTRIBUTIONS

R.M., A.S. and P.P.G. designed and R.M. performed most of the experiments. A.P.F. S.D. R.M. A.S. and B.M.K. designed experiments involving proteomics analysis, and R.M. A.P.F. and S.D. performed the proteomics experiment. J.J.L.L.A. designed the expression and CRISPR knockout constructs together with R.M. R.Q.K. supervised the recombinant protein production. S.Z. and M.G. provided the MOV10KO and WT Huh-7 cells. K.P.K. and G.F. provided recombinant USP18 protein and ISG15 probe precursor constructs. M.K. interpreted the data involving immunoassays. A.S., P.P.G., B.M.K., and A.P.F. supervised the project. R.M., A.P.F., B.M.K., P.P.G., and A.S. interpreted the data and wrote the manuscript with input from all other authors.

---

**Supplementary Fig. 1:**
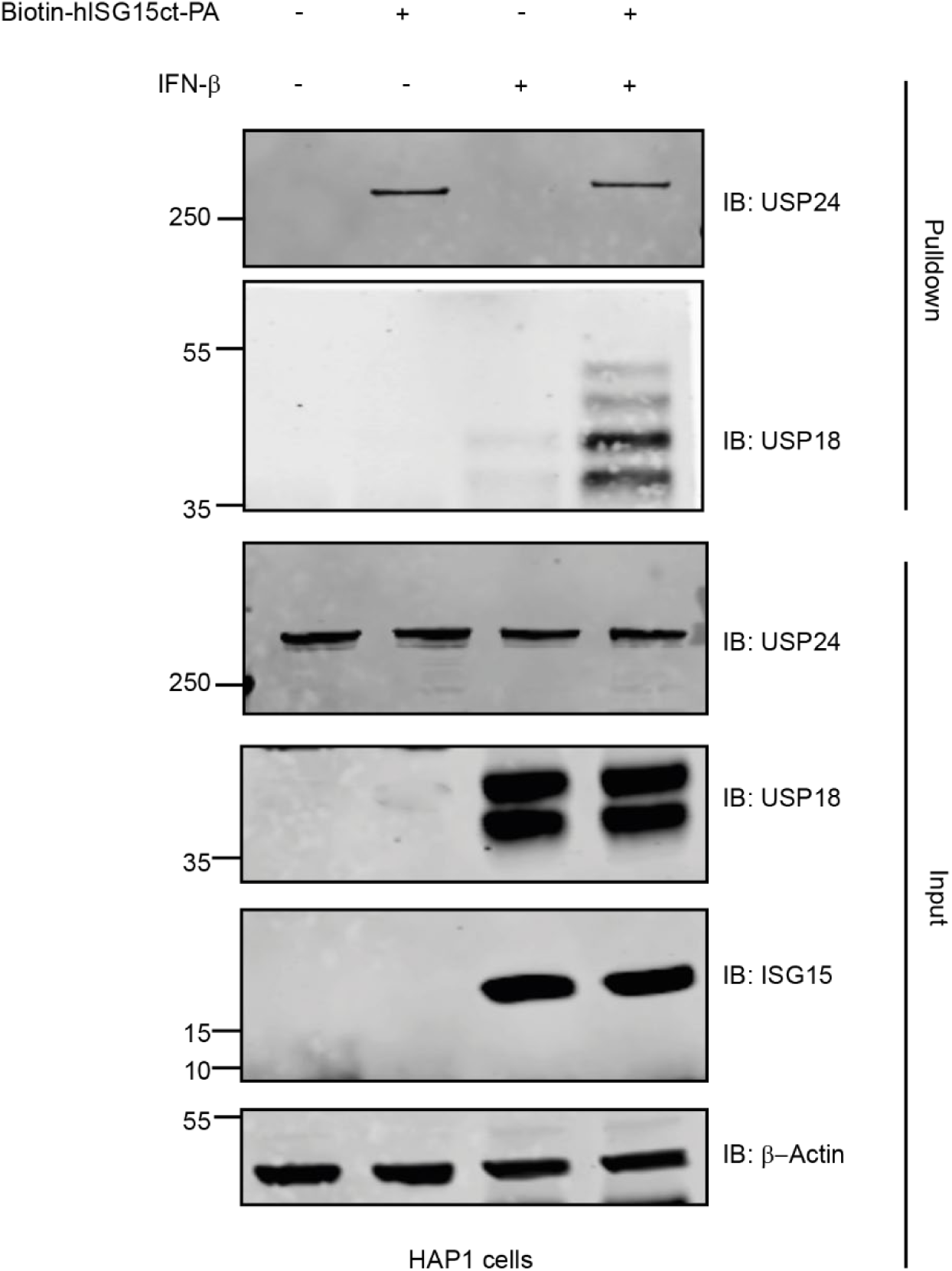
Endogenous USP24 pull-down with Biotin-hISG15_CT_-PA from HAP1 cells. HAP-1 cell lysates from IFN-β stimulated or untreated cells were incubated with 5 µM of Biotin-hISG15_CT_-PA or DMSO for 1 h at 37°C, followed by pulldown with Neutravidin beads and immunoblot analysis against USP24, along with USP18 and ISG15 as positive controls for IFN-β stimulation and β-actin as loading control. Related to Figure 1D.

**Supplementary Fig. 2:**
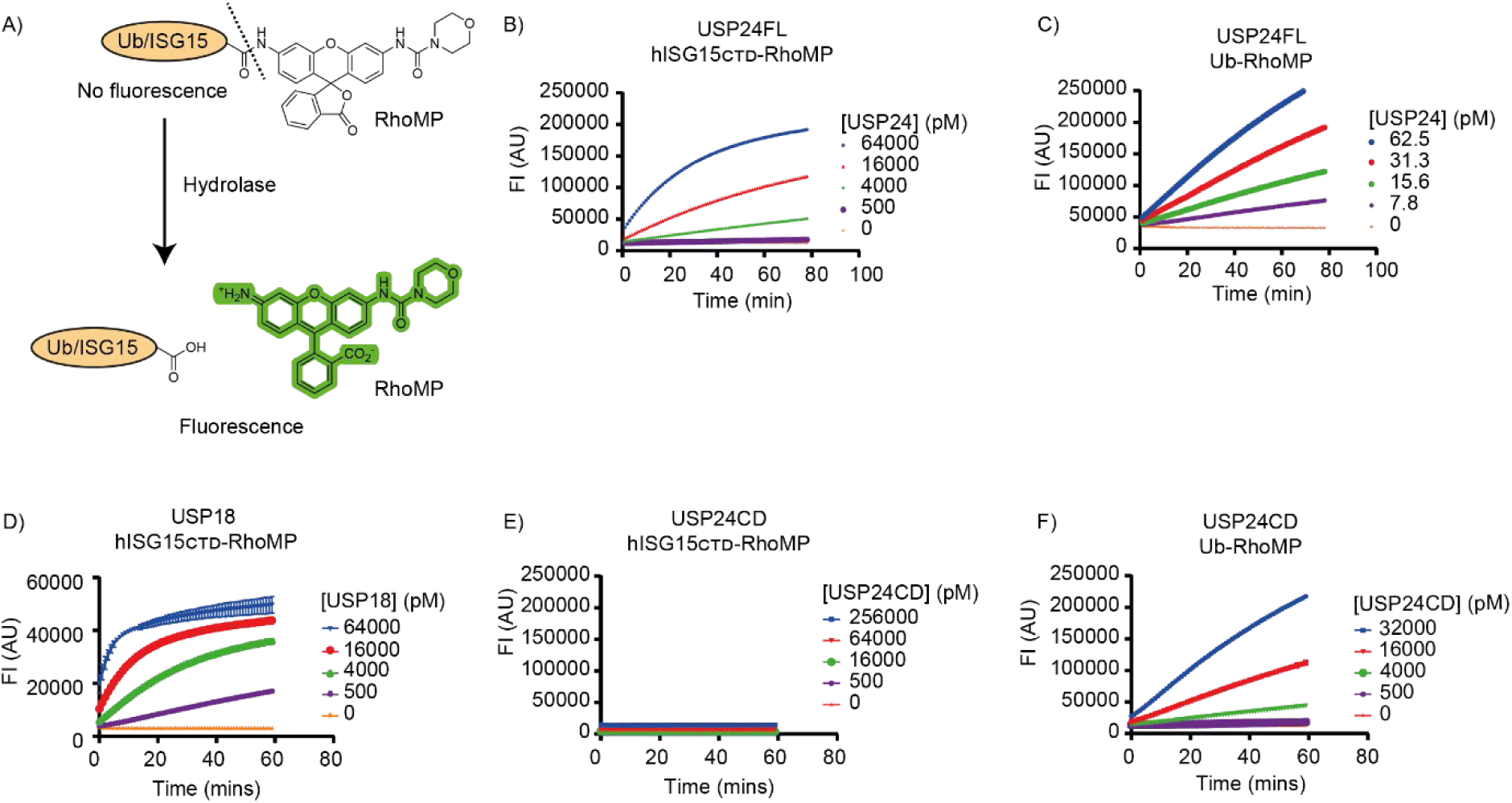
**(A)** Schematic representation of the enzyme activity assay for deconjugating enzymes utilizing a Ub or ISG15 fluorogenic substrate. A fluorophore (Rho-MP) is coupled to the C-terminal carboxylate of synthetic Ub/ISG15, which releases the fluorophore upon processing by a deconjugating enzyme and leads to increased fluorescence intensity over time. The emitted fluorescence intensity is proportional to the enzyme activity. Enzymatic activity measurement of **(B, C)** USP24FL, **(D)** USP18, **(E, F)** USP24CD utilizing Ub/ISG15_CTD_RhoMP assay. The indicated concentration of recombinant enzyme was incubated with 400 nM of the indicated substrates followed by fluorescence intensity measurements for 1 h. Related to Figure 2.

**Supplementary Fig. 3:**
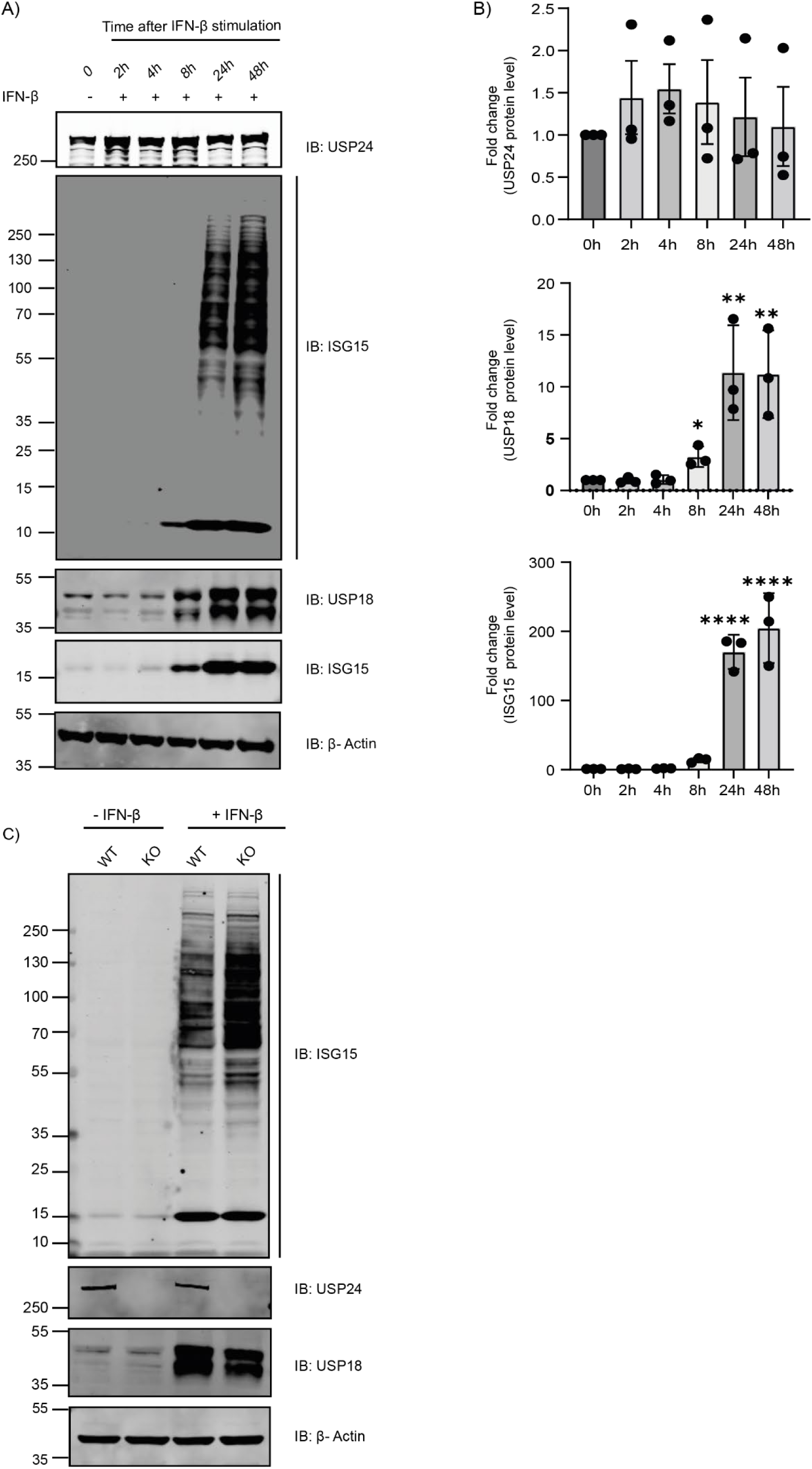
**(A)** Time screening for optimal expression of ISGs and maximum ISG15 conjugate formation after IFNβ treatment. HeLa cells were treated with IFNβ for the indicated time. Immunoblot confirmed the protein levels of USP24, ISG15, and USP18, where β-actin served as loading control. **(B)** Quantification of the fold change of protein levels of USP24, USP18, and ISG15 upon IFNβ stimulation over the indicated time period, n=3 independent experiments. Data is normalized as per negative control. Bar graphs report mean, error bars reflect ± s.d. All significant values were calculated using Student’s t-test: *p < 0.01, **p < 0.001, ****p < 0.0001, NS = not significant. **(C)** Analysis of HeLa USP24 knockout cells for ISGylation profile. Cell lysates from untreated and 48h post-IFNβ treatment were subjected to SDS-PAGE. Immunoblot showed the protein levels of USP24, USP18, and ISG15, confirming USP24 knockout. β-Actin served as loading control. Related to Figure 3A.

**Supplementary Fig. 4:**
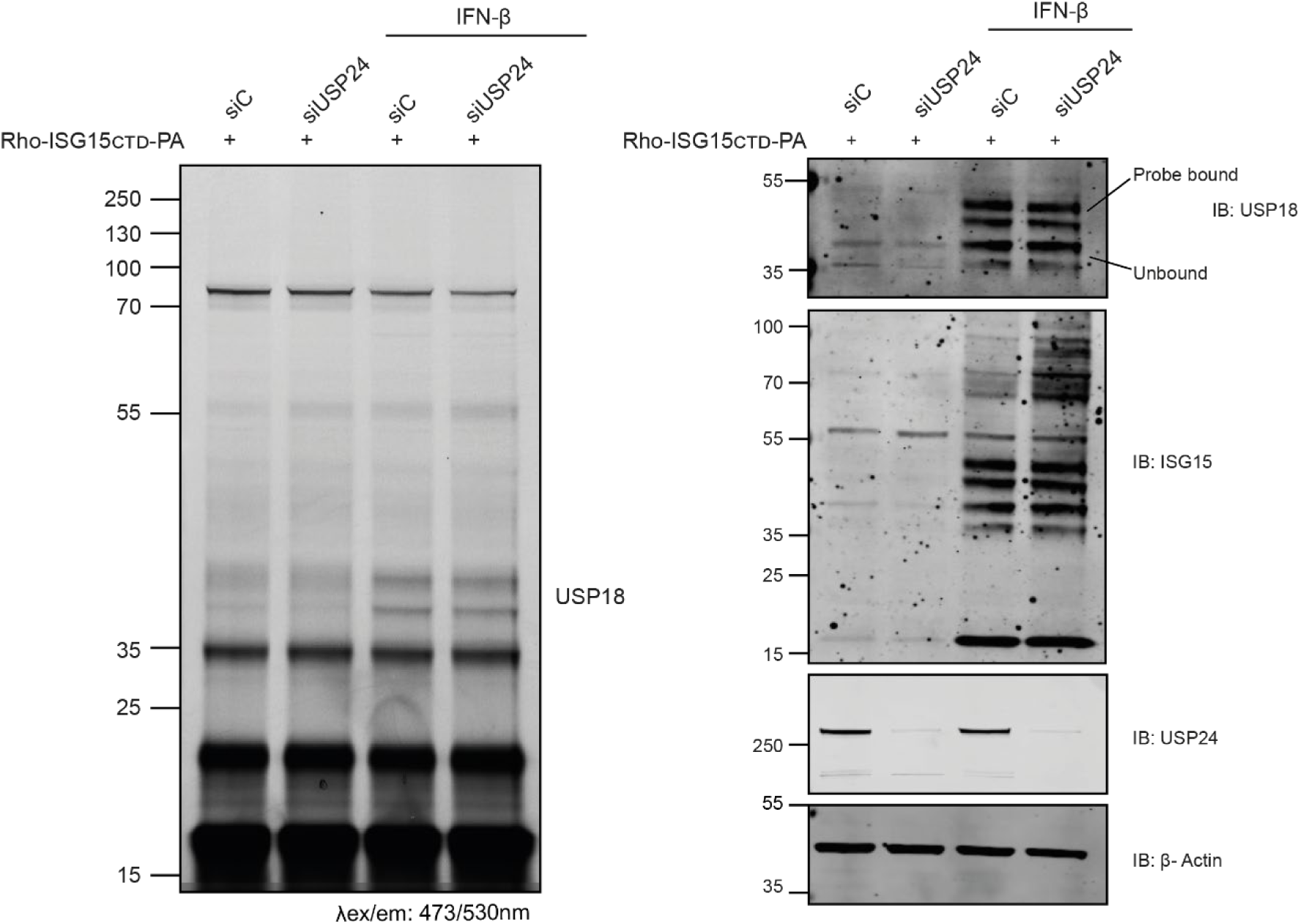
Assessment of enzyme activity of endogenous USP18 in HeLa cell lysate. Cells were treated with either siC or siUSP24, followed by IFN-β stimulation for 48 h. Cells treated with either siC or siUSP24 but not treated with IFN-β were taken along as negative control. Rho-ISG15_CT_-PA probe (1 µM) was incubated with cell lysate for 30 mins at 37°C. Following SDS-PAGE, the gel was scanned for fluorescence at the indicated wavelength (left panel). Immunoblot confirms the protein levels of USP18, ISG15, and USP24, where β-actin served as loading control (right panel). Related to Figure 3.

**Supplementary Fig. 5:**
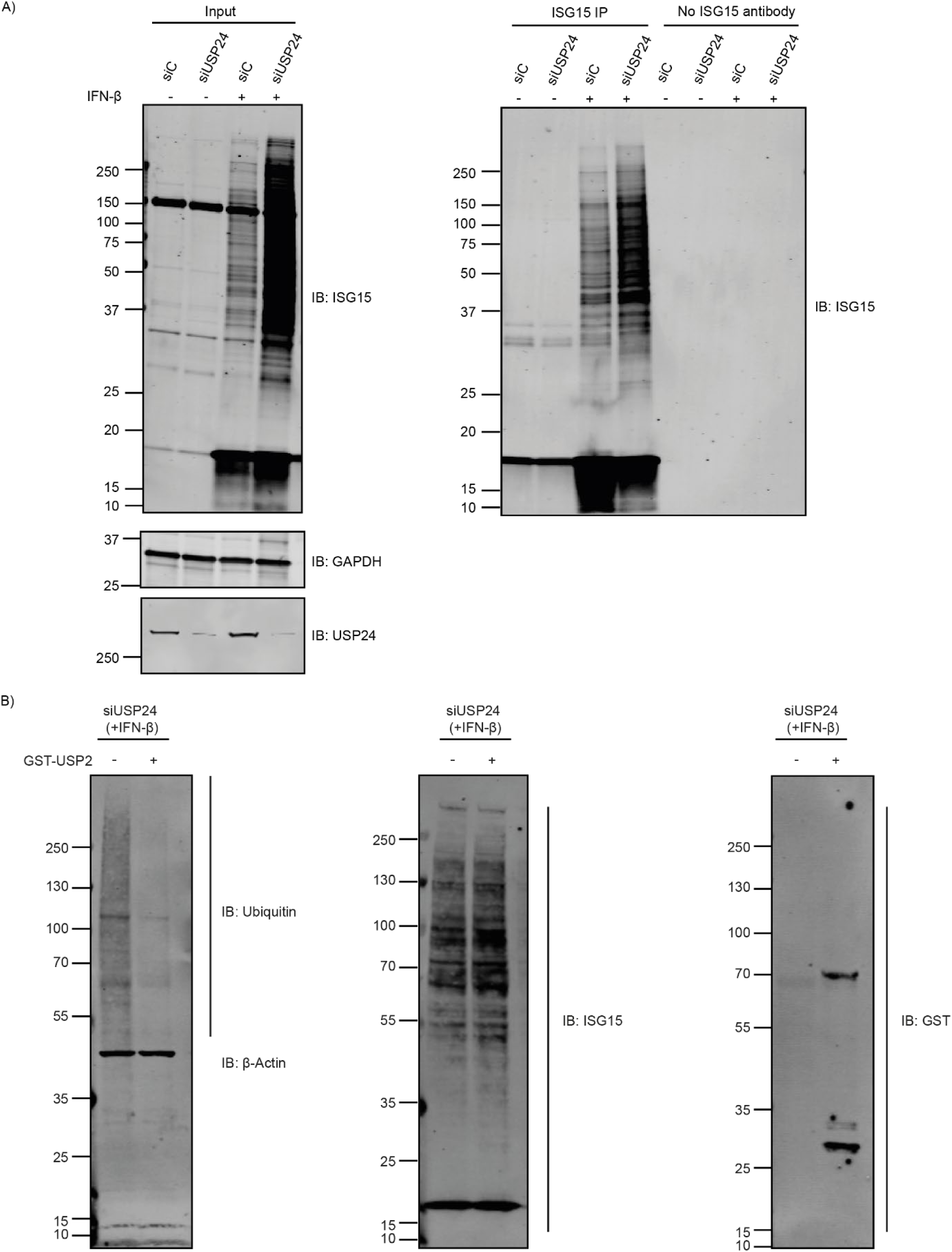
**(A)** ISG15 immunoprecipitation western blot depicting the input samples (left panel) and the immunoprecipitated samples (right panel). As a negative control, samples not treated with ISG15 antibody (beads only) were taken along. Immunoblot confirms protein levels of ISG15 and USP24. GAPDH shows the protein loading control. **(B)** IFN-β treated USP24 depleted HeLa cell lysates were incubated with 1 µM USP2 for 1h at 37°C. As a control, cell lysates not treated with USP2 were taken along. Western blot shows the protein level of GST-USP2 in the lysate, where ubiquitin and ISG15-specific blots highlight the effect of USP2 on the Ubiquitome and ISGylome respectively. β-Actin served as loading control. Related to Figure 4.

**Supplementary Fig. 6:**
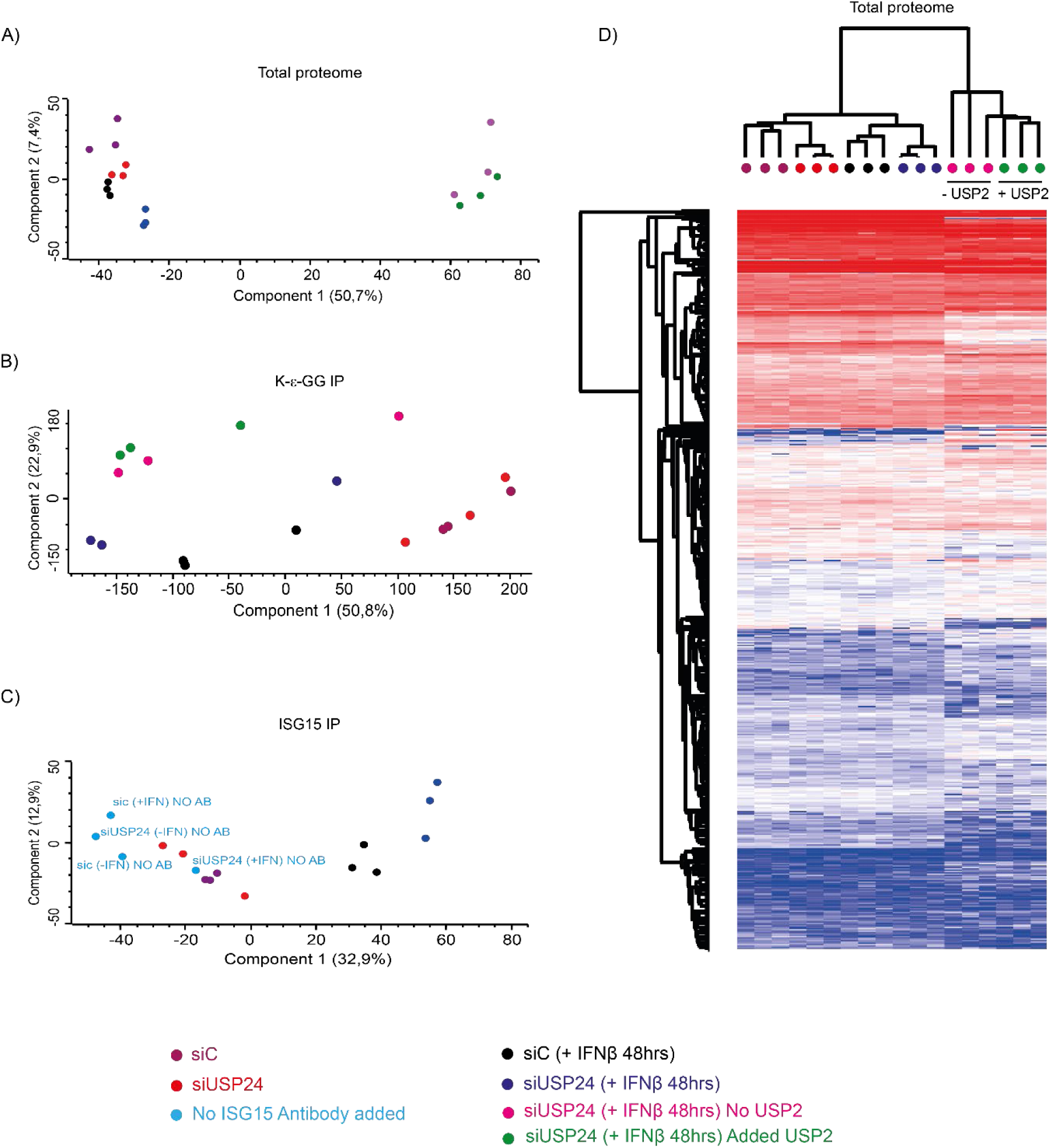
Principal component analysis of the **(A)** total proteome, **(B)** GG-peptidome, and **(C)** ISG15 interactome data-independent acquisition (DIA) data using components with the highest explained variance, n = 3. Colored dots indicated represent cell conditions. **(D)** Heat map showing the hierarchical clustering of the significantly affected genes (p-value < 0.05) from the Total proteome data set. Clustering was done based on Euclidean distance (pre-processed with k-means, 300 clusters, 1000 iterations), where each column in the heatmap is a sample and each row is a protein. The LFQ intensity values were normalized by Z score. Source data are provided as a Source Data file: S1. Related to Figure 4.

**Supplementary figure 7:**
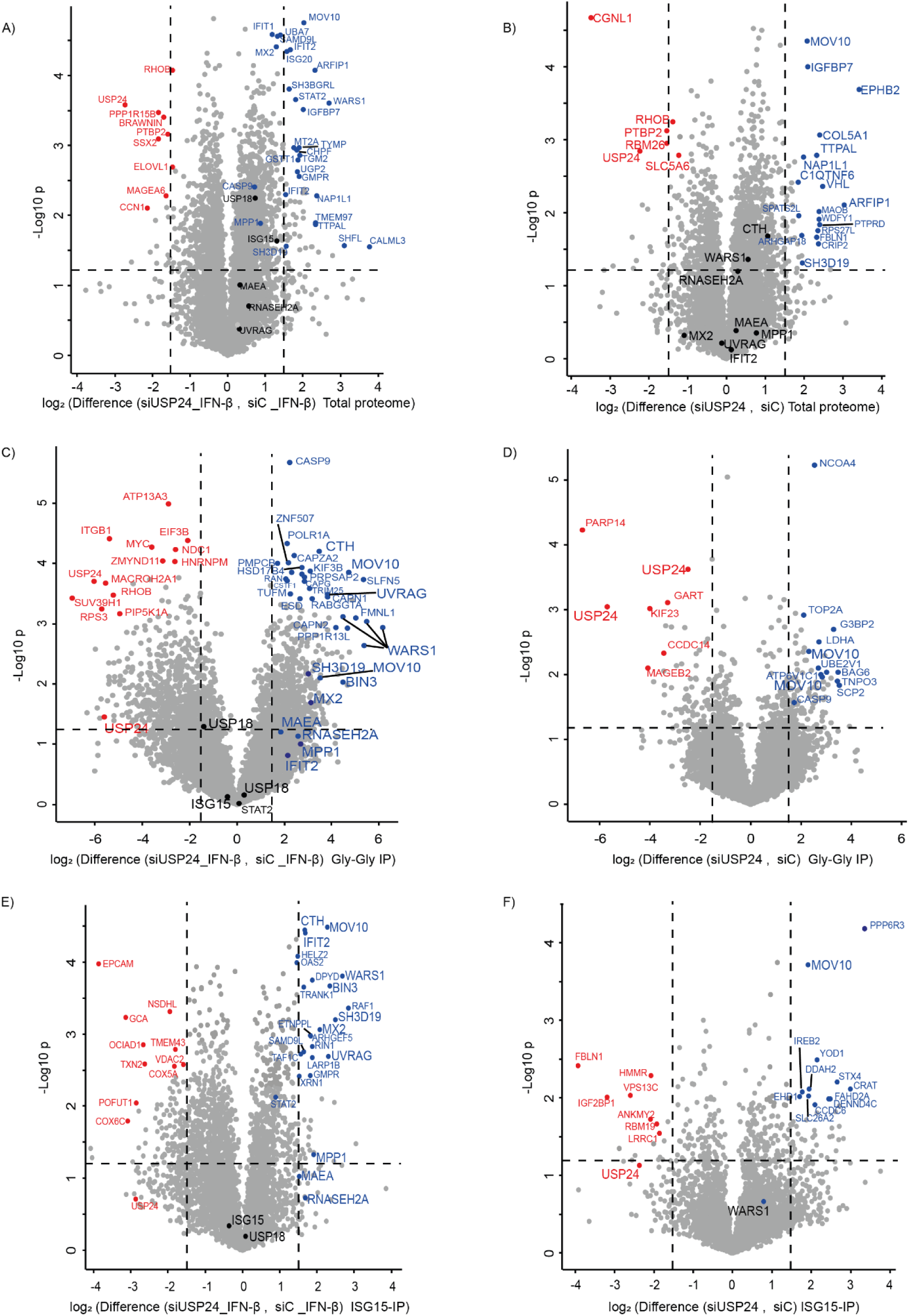
**(A, B)** Volcano plots of the total proteome analysis of HeLa cells with/without USP24 knockdown (-/+ IFN-β, 1:1000, 48 h). Dashed lines indicate significance cutoff at a p-value of 0.05 (-log10 = 1.3) in y-axis and log2 ≥ 1.5 cutoff at fold change in x-axis, n=3 independent experiments. All the proteins identified that are enriched with USP24 depletion are highlighted in blue, while the proteins downregulated by USP24 depletion are highlighted in red. ISG15 and USP18 showed no significant changes. *Left volcano plot (A)*: IFN-β stimulated cells. *Right volcano plot (B)*: HeLa cells not treated with IFN-β (Negative control). Source data are provided as a Supplementary Data S1. **(C, D)** di-Gly immunoprecipitation in HeLa WT cells and USP24 knockdown HeLa cells (-/+ IFN-β, 1:1000, 48 h). Dashed lines indicate significance cutoff at a p-value of 0.05 (-log10 = 1.3), n=3 independent experiments. All the proteins identified that are upregulated with USP24 depletion and enriched by di-Gly immunoprecipitation are highlighted in blue, the proteins downregulated by USP24 depletion are highlighted in red. ISG15 and USP18 showed no significant changes. *Left volcano plot (C)*: IFN-β stimulated cells. *Right volcano plot (D)*: HeLa cells not treated with IFN-β (Negative control). Source data are provided as Supplementary Data S2. **(E, F)** ISGylome in HeLa WT cells and USP24 knockdown HeLa cells (-/+ IFN-β, 1:1000, 48 h). Dashed lines indicate significance cutoff at a p-value of 0.05 (-log10 = 1.3), n=3 independent experiments. All the proteins identified that are upregulated by USP24 depletion are highlighted in blue, while the proteins downregulated by USP24 depletion are highlighted in red. ISG15 and USP18 showed no significant changes. *Left volcano plot (E)*: IFN-β stimulated cells. *Right volcano plot (F)*: HeLa cells not treated with IFN-β (Negative control). Source data are provided as Supplementary Data S3. Related to Figure 4.

**Supplementary Fig. 8:**
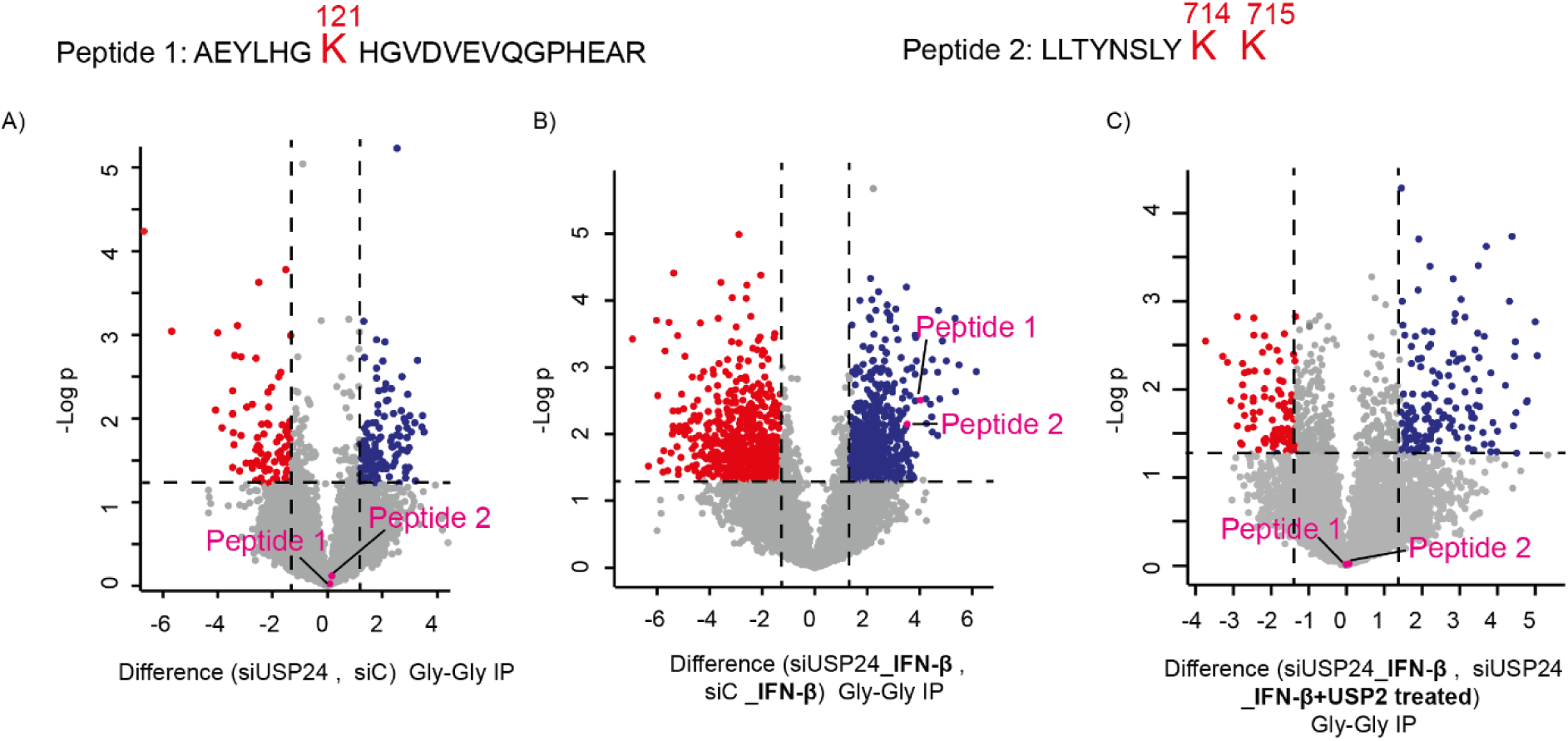
Analysis of the ISG15 modification site on MOV10. Two tryptic peptides of MOV10 were identified to be ISG15 conjugated: Peptide 1 and Peptide 2. (A) Peptide 1 and Peptide 2 were not enriched by Gly-Gly antibody in IFN-β untreated cells. (B) Peptide 1 and Peptide 2 were enriched by Gly-Gly antibody in IFN-β treated cells. (C) Peptide 1 and Peptide 2 showed no significant processing by USP2 in IFN-β treated cell lysates. The peptides revealed two unique Lysines (K121 and K714) that can be ISG15 modified. An additional lysine residue (K715) just beside K714 is also considered due to their close proximity. Lysine residues identified to be ISG15 modified are highlighted in red color.

**Supplementary Table 1:**
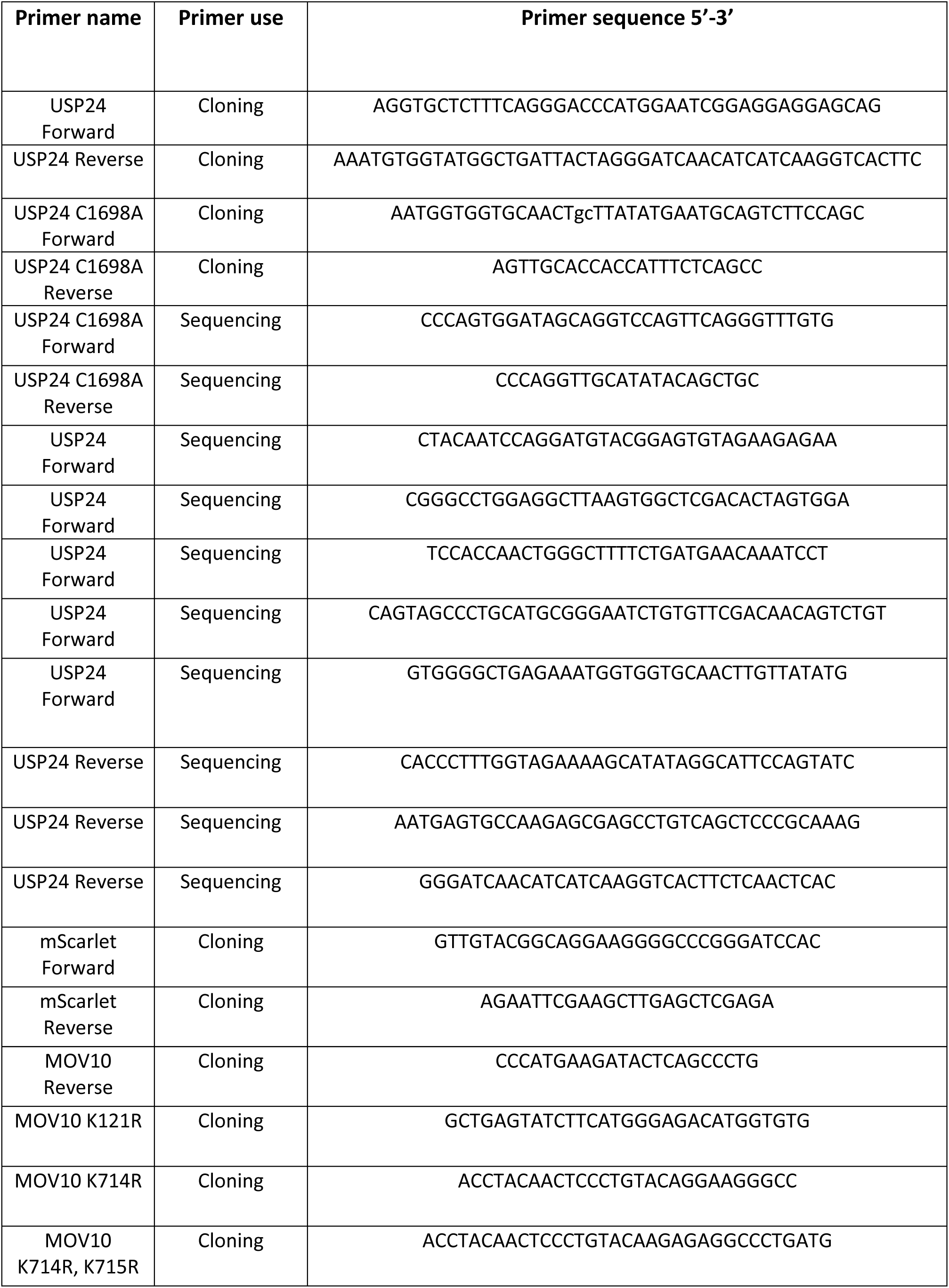
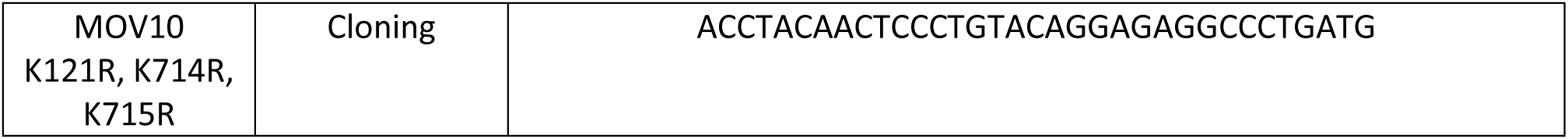
Details of the primers used in the study.

## REFERENCES

1 Lee, J. M., Hammaren, H. M., Savitski, M. M. & Baek, S. H. Control of protein stability by post-translational modifications. Nat Commun 14, 201, doi:10.1038/s41467-023-35795-8 (2023).

2 Kang, J. A., Kim, Y. J. & Jeon, Y. J. The diverse repertoire of ISG15: more intricate than initially thought. Exp Mol Med 54, 1779–1792, doi:10.1038/s12276-022-00872-3 (2022).

3 Wang, Y. E., Pernet, O. & Lee, B. Regulation of the nucleocytoplasmic trafficking of viral and cellular proteins by ubiquitin and small ubiquitin-related modifiers. Biol Cell 104, 121–138, doi:10.1111/boc.201100105 (2012).

4 Zhong, Q. et al. Protein posttranslational modifications in health and diseases: Functions, regulatory mechanisms, and therapeutic implications. MedComm (2020) 4, e261, doi:10.1002/mco2.261 (2023).

5 Narasimhan, J. et al. Crystal structure of the interferon-induced ubiquitin-like protein ISG15. J Biol Chem 280, 27356–27365, doi:10.1074/jbc.M502814200 (2005).

6 Perng, Y. C. & Lenschow, D. J. ISG15 in antiviral immunity and beyond. Nat Rev Microbiol 16, 423–439, doi:10.1038/s41579-018-0020-5 (2018).

7 Pitha-Rowe, I., Hassel, B. A. & Dmitrovsky, E. Involvement of UBE1L in ISG15 conjugation during retinoid-induced differentiation of acute promyelocytic leukemia. J Biol Chem 279, 18178–18187, doi:10.1074/jbc.M309259200 (2004).

8 Orfali, N. et al. Inhibition of UBE2L6 attenuates ISGylation and impedes ATRA-induced differentiation of leukemic cells. Mol Oncol 14, 1297–1309, doi:10.1002/1878-0261.12614 (2020).

9 Wong, J. J., Pung, Y. F., Sze, N. S. & Chin, K. C. HERC5 is an IFN-induced HECT-type E3 protein ligase that mediates type I IFN-induced ISGylation of protein targets. Proc Natl Acad Sci U S A 103, 10735–10740, doi:10.1073/pnas.0600397103 (2006).

10 Wu, S. F. et al. RIG-I regulates myeloid differentiation by promoting TRIM25-mediated ISGylation. Proc Natl Acad Sci U S A 117, 14395–14404, doi:10.1073/pnas.1918596117 (2020).

11 Xiong, T. C. et al. The E3 ubiquitin ligase ARIH1 promotes antiviral immunity and autoimmunity by inducing mono-ISGylation and oligomerization of cGAS. Nat Commun 13, 5973, doi:10.1038/s41467-022-33671-5 (2022).

12 Durfee, L. A., Lyon, N., Seo, K. & Huibregtse, J. M. The ISG15 conjugation system broadly targets newly synthesized proteins: implications for the antiviral function of ISG15. Mol Cell 38, 722–732, doi:10.1016/j.molcel.2010.05.002 (2010).

13 Basters, A. et al. Structural basis of the specificity of USP18 toward ISG15. Nat Struct Mol Biol 24, 270–278, doi:10.1038/nsmb.3371 (2017).

14 Basters, A. et al. Molecular characterization of ubiquitin-specific protease 18 reveals substrate specificity for interferon-stimulated gene 15. FEBS J 281, 1918–1928, doi:10.1111/febs.12754 (2014).

15 Malakhov, M. P., Malakhova, O. A., Kim, K. I., Ritchie, K. J. & Zhang, D. E. UBP43 (USP18) specifically removes ISG15 from conjugated proteins. J Biol Chem 277, 9976–9981, doi:10.1074/jbc.M109078200 (2002).

16 Serniwka, S. A. & Shaw, G. S. The structure of the UbcH8-ubiquitin complex shows a unique ubiquitin interaction site. Biochemistry 48, 12169–12179, doi:10.1021/bi901686j (2009).

17 Catic, A. et al. Screen for ISG15-crossreactive deubiquitinases. PLoS One 2, e679, doi:10.1371/journal.pone.0000679 (2007).

18 Ye, Y. et al. Polyubiquitin binding and cross-reactivity in the USP domain deubiquitinase USP21. EMBO Rep 12, 350–357, doi:10.1038/embor.2011.17 (2011).

19 Hemelaar, J. et al. Chemistry-based functional proteomics: mechanism-based activity-profiling tools for ubiquitin and ubiquitin-like specific proteases. J Proteome Res 3, 268–276, doi:10.1021/pr0341080 (2004).

20 Gan, J. et al. USP16 is an ISG15 cross-reactive deubiquitinase that targets pro-ISG15 and ISGylated proteins involved in metabolism. Proc Natl Acad Sci U S A 120, e2315163120, doi:10.1073/pnas.2315163120 (2023).

21 O’Dea, R. et al. Molecular basis for ubiquitin/Fubi cross-reactivity in USP16 and USP36. Nat Chem Biol 19, 1394–1405, doi:10.1038/s41589-023-01388-1 (2023).

22 Wang, Y. C. et al. USP24 induces IL-6 in tumor-associated microenvironment by stabilizing p300 and beta-TrCP and promotes cancer malignancy. Nat Commun 9, 3996, doi:10.1038/s41467-018-06178-1 (2018).

23 Zhang, L. et al. The deubiquitinating enzyme USP24 is a regulator of the UV damage response. Cell Rep 10, 140–147, doi:10.1016/j.celrep.2014.12.024 (2015).

24 Nawaz, A., Shilikbay, T., Skariah, G. & Ceman, S. Unwinding the roles of RNA helicase MOV10. Wiley Interdiscip Rev RNA 13, e1682, doi:10.1002/wrna.1682 (2022).

25 Cuevas, R. A. et al. MOV10 Provides Antiviral Activity against RNA Viruses by Enhancing RIG-I-MAVS-Independent IFN Induction. J Immunol 196, 3877–3886, doi:10.4049/jimmunol.1501359 (2016).

26 Zang, L. et al. Ubiquitin-specific protease 24 promotes EV71 infection by restricting K63-linked polyubiquitination of TBK1. Virol Sin 38, 75–83, doi:10.1016/j.virs.2022.11.001 (2023).

27 Geurink, P. P. et al. Profiling DUBs and Ubl-specific proteases with activity-based probes. Methods Enzymol 618, 357–387, doi:10.1016/bs.mie.2018.12.037 (2019).

28 Kooij, R. et al. Small-Molecule Activity-Based Probe for Monitoring Ubiquitin C-Terminal Hydrolase L1 (UCHL1) Activity in Live Cells and Zebrafish Embryos. J Am Chem Soc 142, 16825–16841, doi:10.1021/jacs.0c07726 (2020).

29 Xin, B. T. et al. Total chemical synthesis of murine ISG15 and an activity-based probe with physiological binding properties. Org Biomol Chem 17, 10148–10152, doi:10.1039/c9ob02127b (2019).

30 Potter, J. L., Narasimhan, J., Mende-Mueller, L. & Haas, A. L. Precursor processing of pro-ISG15/UCRP, an interferon-beta-induced ubiquitin-like protein. J Biol Chem 274, 25061–25068, doi:10.1074/jbc.274.35.25061 (1999).

31 McNab, F., Mayer-Barber, K., Sher, A., Wack, A. & O’Garra, A. Type I interferons in infectious disease. Nat Rev Immunol 15, 87–103, doi:10.1038/nri3787 (2015).

32 Gregersen, L. H. et al. MOV10 Is a 5’ to 3’ RNA helicase contributing to UPF1 mRNA target degradation by translocation along 3’ UTRs. Mol Cell 54, 573–585, doi:10.1016/j.molcel.2014.03.017 (2014).

33 Reyes-Turcu, F. E., Ventii, K. H. & Wilkinson, K. D. Regulation and cellular roles of ubiquitin-specific deubiquitinating enzymes. Annu Rev Biochem 78, 363–397, doi:10.1146/annurev.biochem.78.082307.091526 (2009).

34 Zhang, Q., Jia, Q., Gao, W. & Zhang, W. The Role of Deubiquitinases in Virus Replication and Host Innate Immune Response. Front Microbiol 13, 839624, doi:10.3389/fmicb.2022.839624 (2022).

35 Wada, H., Kito, K., Caskey, L. S., Yeh, E. T. & Kamitani, T. Cleavage of the C-terminus of NEDD8 by UCH-L3. Biochem Biophys Res Commun 251, 688–692, doi:10.1006/bbrc.1998.9532 (1998).

36 Munnur, D., Banducci-Karp, A. & Sanyal, S. ISG15 driven cellular responses to virus infection. Biochem Soc Trans 50, 1837–1846, doi:10.1042/BST20220839 (2022).

37 Song, X. et al. Related cellular signaling and consequent pathophysiological outcomes of ubiquitin specific protease 24. Life Sci 342, 122512, doi:10.1016/j.lfs.2024.122512 (2024).

38 Ketscher, L. et al. Selective inactivation of USP18 isopeptidase activity in vivo enhances ISG15 conjugation and viral resistance. Proc Natl Acad Sci U S A 112, 1577–1582, doi:10.1073/pnas.1412881112 (2015).

39 Arimoto, K. I. et al. STAT2 is an essential adaptor in USP18-mediated suppression of type I interferon signaling. Nat Struct Mol Biol 24, 279–289, doi:10.1038/nsmb.3378 (2017).

40 Zhao, X. et al. Cellular targets and lysine selectivity of the HERC5 ISG15 ligase. iScience 27, 108820, doi:10.1016/j.isci.2024.108820 (2024).

41 Song, Z. W. et al. Altered mRNA levels of MOV10, A3G, and IFN-alpha in patients with chronic hepatitis B. J Microbiol 52, 510–514, doi:10.1007/s12275-014-3467-8 (2014).

42 Kali, S. K., Droge, P. & Murugan, P. Interferon beta, an enhancer of the innate immune response against SARS-CoV-2 infection. Microb Pathog 158, 105105, doi:10.1016/j.micpath.2021.105105 (2021).

43 Huang, C. T. et al. Enhancement of the IFN-beta-induced host signature informs repurposed drugs for COVID-19. Heliyon 6, e05646, doi:10.1016/j.heliyon.2020.e05646 (2020).

44 Borden, E. C. Interferons alpha and beta in cancer: therapeutic opportunities from new insights. Nat Rev Drug Discov 18, 219–234, doi:10.1038/s41573-018-0011-2 (2019).

45 Wang, S. A. et al. NCI677397 targeting USP24-mediated induction of lipid peroxidation induces ferroptosis in drug-resistant cancer cells. Mol Oncol, doi:10.1002/1878-0261.13574 (2023).

46 Häusler, L. The role of the deubiquitinase USP24 in multiple myeloma (2017)

47 Garcia-Nafria, J., Watson, J. F. & Greger, I. H. IVA cloning: A single-tube universal cloning system exploiting bacterial In Vivo Assembly. Sci Rep 6, 27459, doi:10.1038/srep27459 (2016).

48 Wang, L., Sola, I., Enjuanes, L. & Zuniga, S. MOV10 Helicase Interacts with Coronavirus Nucleocapsid Protein and Has Antiviral Activity. mBio 12, e0131621, doi:10.1128/mBio.01316-21 (2021).

49 Rocha, N. et al. Cholesterol sensor ORP1L contacts the ER protein VAP to control Rab7-RILP-p150 Glued and late endosome positioning. J Cell Biol 185, 1209–1225, doi:10.1083/jcb.200811005 (2009).

50 Pinto-Fernandez, A. et al. Comprehensive Landscape of Active Deubiquitinating Enzymes Profiled by Advanced Chemoproteomics. Front Chem 7, 592, doi:10.3389/fchem.2019.00592 (2019).

51 Kelley, L. A., Mezulis, S., Yates, C. M., Wass, M. N. & Sternberg, M. J. The Phyre2 web portal for protein modeling, prediction and analysis. Nat Protoc 10, 845–858, doi:10.1038/nprot.2015.053 (2015).

52 Waterhouse, A. et al. SWISS-MODEL: homology modelling of protein structures and complexes. Nucleic Acids Res 46, W296–W303, doi:10.1093/nar/gky427 (2018).

53 Ozen, A. et al. Selectively Modulating Conformational States of USP7 Catalytic Domain for Activation. Structure 26, 72–84 e77, doi:10.1016/j.str.2017.11.010 (2018).

54 Meier, F. et al. diaPASEF: parallel accumulation-serial fragmentation combined with data-independent acquisition. Nat Methods 17, 1229–1236, doi:10.1038/s41592-020-00998-0 (2020).

55 Skowronek, P. et al. Rapid and In-Depth Coverage of the (Phospho-)Proteome With Deep Libraries and Optimal Window Design for dia-PASEF. Mol Cell Proteomics 21, 100279, doi:10.1016/j.mcpro.2022.100279 (2022).

56 Tyanova, S. et al. The Perseus computational platform for comprehensive analysis of (prote)omics data. Nat Methods 13, 731–740, doi:10.1038/nmeth.3901 (2016).

57 Cline, M. S. et al. Integration of biological networks and gene expression data using Cytoscape. Nat Protoc 2, 2366–2382, doi:10.1038/nprot.2007.324 (2007).

